# Genetic association models are robust to common population kinship estimation biases

**DOI:** 10.1101/2022.11.07.515490

**Authors:** Zhuoran Hou, Alejandro Ochoa

## Abstract

Common genetic association models for structured populations, including Principal Component Analysis (PCA) and Linear Mixed-effects Models (LMM), model the correlation structure between individuals using population kinship matrices, also known as Genetic Relatedness Matrices or “GRMs”. However, the most common kinship estimators can have severe biases that were only recently determined. Here we characterize the effect of these kinship biases on genetic association. We employ a large simulated admixed family and genotypes from the 1000 Genomes Project, both with simulated traits, to evaluate key kinship estimators. Remarkably, we find practically invariant association statistics for kinship matrices of different bias types (matching all other features). We then prove using statistical theory and linear algebra that LMM association tests are invariant to these kinship biases, and PCA approximately so. Our proof shows that the intercept and relatedness effect coefficients compensate for the kinship bias, an argument that extends to generalized linear models. As a corollary, association testing is also invariant to changing the reference ancestral population of the kinship matrix. Lastly, we observed that all kinship estimators, except for popkin ROM, can give improper non-positive semidefinite matrices, which can be problematic although some LMMs handle them surprisingly well, and condition numbers can be used to choose kinship estimators. Overall, we find that existing association studies are robust to kinship estimation bias, and our calculations may help improve association methods by taking advantage of this unexpected robustness, as well as help determine the effects of kinship bias in related problems.

## 1 Introduction

The goal of genetic association is to detect loci that are related to a specific trait, either causally or by proximity to causal loci. When applied to structured populations with admixed individuals, multiethnic cohorts, or close relatives, controlling for relatedness is crucial to avoid spurious associations and loss of power (Devlin and Roeder, 1999; Voight and Pritchard, 2005; Astle and Balding, 2009; Yao and Ochoa, 2022). The most popular association models for structured populations are Linear Mixed-effects Models (LMM) and Principal Component Analysis (PCA), which are closely related except LMM is capable of modeling high-dimensional structures whereas PCA is strictly a low-dimensional model (Astle and Balding, 2009; Hoffman, 2013; Yao and Ochoa, 2022).

Various association models, including both PCA and LMM, parametrize relatedness using kinship matrices, also known as Genetic Relatedness Matrices or “GRMs”. Kinship coefficients are well suited for this task since they model the covariance structure of genotypes (Malécot, 1948; Jacquard, 1970). Kinship is often encountered in family studies, where they reflect recent relatedness and can be calculated from pedigrees (Wright, 1922; Emik and Terrill, 1949; García-Cortés, 2015). However, as kinship is defined as a probability of identity by descent, it may also capture ancient population relatedness (Malécot, 1948; Astle and Balding, 2009), and common non-parametric kinship estimators from genotypes indeed include population structure in their estimates (Ochoa and Storey, 2021). In LMMs, the kinship matrix is an explicit parameter determining the random effect covariance structure (Xie et al., 1998; Yu et al., 2006; Aulchenko et al., 2007; Astle and Balding, 2009; Kang et al., 2008; Kang et al., 2010; Zhou and Stephens, 2012; Yang et al., 2014; Loh et al., 2015; Sul et al., 2018). In PCA, the principal components (PCs) are in practice the eigenvectors of an empirical genetic covariance matrix that is equivalent to the most common kinship estimator (Price et al., 2006; Astle and Balding, 2009; Hoffman, 2013; Yao and Ochoa, 2022).

Although several kinship estimators have been used with LMMs in the past, work from the last 15 years has converged on what we call the “standard” kinship estimator, which is the same estimator used in PCA and other related models (Price et al., 2006; Astle and Balding, 2009; Rakovski and Stram, 2009; Thornton and McPeek, 2010; Yang et al., 2010; Yang et al., 2011; Zhou and Stephens, 2012; Speed et al., 2012; Yang et al., 2014; Speed and Balding, 2015; Loh et al., 2015; Wang et al., 2017; Sul et al., 2018). The impetus of our work is the recent characterization of a complex bias for this standard estimator, which varies for every pair of individuals (Weir and Goudet, 2017; Ochoa and Storey, 2021). These recent works also produced two new kinship estimators, which we are interested in characterizing in the context of association. The Weir-Goudet (WG) estimator constitutes a key improvement in that it has a uniformly downward bias (Weir and Goudet, 2017; Ochoa and Storey, 2021). Lastly, the popkin estimator is the only unbiased estimator under arbitrary relatedness (Ochoa and Storey, 2021). To the best of our knowledge, the new WG and popkin estimators have not been used in association studies before, but represent potential improvements over the use of the standard estimator for association.

One potential confounder when comparing the above kinship estimators is that the standard estimator upweighs rare variants in a formulation previously called “mean-of-ratios” (MOR), whereas WG and popkin do not, instead following a “ratio-of-means” (ROM) estimation strategy (Bhatia et al., 2013; Ochoa and Storey, 2021). Recent work also formulated a ROM version of the standard estimator, which has a more predictable bias than the widely used MOR version (Ochoa and Storey, 2021). Following a locus weight formulation that allows the standard estimator to weigh loci in both ways (Wang et al., 2017), here we generalize the popkin and WG estimators to have both MOR and ROM versions, to test estimators without confounding by locus weighing strategy.

In this work, we originally hypothesized that kinship estimation bias would affect association testing. We perform evaluations using an admixed family simulation (Yao and Ochoa, 2022) as well as real genotypes from the 1000 Genomes project (Consortium, 2010; 1000 Genomes Project Consortium et al., 2012; Fairley et al., 2020), in both cases with simulated traits, to characterize type I error control and power using robust statistics. Surprisingly, we find that both LMM and PCA association statistics are largely invariant to kinship estimation bias. We theoretically characterize the conditions under which these kinship biases result in invariant association statistics, which encompass changing ancestral population in the kinship matrix too. As we discover that most kinship estimates are non-positive semidefinite (non-PSD), breaking a key modeling assumption, we perform additional empirical validations and discover that some LMMs can handle these improper covariance matrices surprisingly well. Overall, we find that long-used association approaches are unaffected by the most common kinship estimation biases, and develop theory that may help improve association and related approaches such as heritability estimation.

## 2 Methods

### 2.1 Genetic model

The following genetic model justifies the use of kinship matrices in association studies, and is the basis of all kinship estimation bias calculations that our theoretical work depends upon.

Suppose there are *m* biallelic loci and *n* diploid individuals. The genotype *x*_*ij*_ ∈ {0, 1, 2} at a locus *i* of individual *j* is encoded as the number of reference alleles, for a preselected but otherwise arbitrary reference allele per locus. Genotypes are treated as random variables structured according to relatedness. If *T* is the ancestral population on which allele frequencies are conditioned, 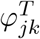 is the kinship coefficient of two individuals *j* and *k*, and 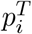 is the ancestral allele frequency at locus *i*, then under the kinship model (Malécot, 1948; Wright, 1949; Jacquard, 1970; Astle and Balding, 2009; Ochoa and Storey, 2021) the expectation and covariance are given by

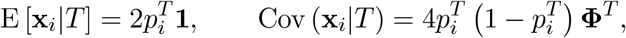

where **x**_*i*_ = (*x*_*ij*_) is the the length-*n* column vector of genotypes at locus *i*, 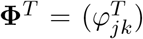 is the *n* × *n* kinship matrix, and **1** is a length-*n* column vector of ones. Both **Φ**^*T*^ and 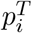 are parameters that depend on the choice of ancestral population, for which the Most Recent Common Ancestor (MRCA) population is the most sensible choice (Ochoa and Storey, 2021). However, one of the results of this work is proof that the choice of ancestral population does not affect association testing.

### 2.2 Kinship estimators

Each subsection below corresponds to a kinship estimator bias type: Popkin is unbiased, while Standard and WG have different bias functions (defined shortly). Each estimator bias type has two locus weight types called *ratio-of-means* (ROM) and *mean-of-ratios* (MOR), a terminology that follows previous convention for these and related estimators (Bhatia et al., 2013; Ochoa and Storey, 2021). Only ROM estimators have closed-form limits. Below 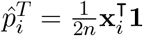 is the standard ancestral allele frequency estimator, where the ⊤ superscript denotes matrix transposition (do not confuse with ancestral population superscript *T*), and 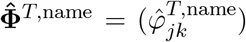 relates the scalar and matrix formulas of each named kinship estimator.

#### 2.2.1 Popkin estimator

The popkin (population kinship) estimator (Ochoa and Storey, 2021), generalized here to include locus weights *w*_*i*_, is given by

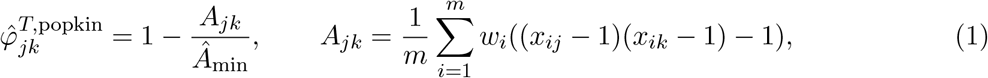

where in this work *Â*_min_ = min_*j≠k*_ *A*_*jk*_, and *w*_*i*_ must be positive but need not add to 1. We consider two broad forms for this estimator. The original ROM estimator has *w*_*i*_ = 1 and has an unbiased almost sure limit as the number of loci *m* go to infinity,

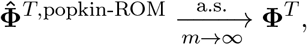

under the assumption that the true minimum kinship is zero. The MOR version, introduced here, upweighs rare variants by using 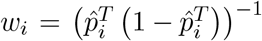; although it has no closed-form limit, it is approximately unbiased as well (Appendix A) and it is connected to the most common estimator, Standard MOR (Appendix B). The use of locus weights here is inspired by previous calculations relating the standard ROM and MOR estimators (Wang et al., 2017).

#### 2.2.2 Standard estimator

The ROM and MOR versions of the standard kinship estimator are, respectively,

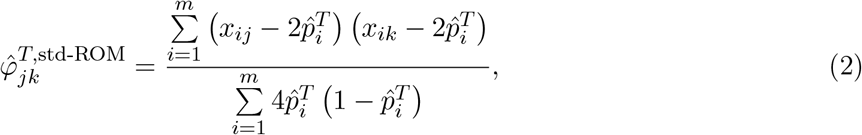

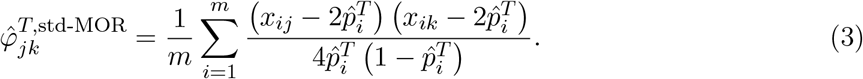

The ROM estimator has a biased limit, which is a function of the true kinship matrix (Ochoa and Storey, 2021):

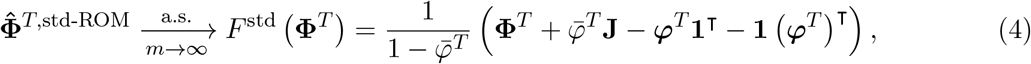

where **J** = **11**^⊤^ is the *n* × *n* matrix of ones, 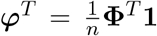 is a length-*n* vector of per-row mean kinship values, and 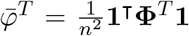 is the scalar overall mean kinship. The MOR estimator does not have closed-form limit, but it is well approximated by Eq. (4) in practice, especially when loci with small minor allele frequencies are excluded prior to calculating this estimate. In Appendix B we prove that, when there are no missing genotypes, the two standard estimators are functions of the corresponding popkin estimators, given by the bias function *F* ^std^:

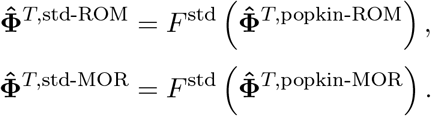

#### 2.2.3 Weir-Goudet estimator

The ROM version of the Weir-Goudet (WG) kinship estimator is given by (Weir and Goudet, 2017; Ochoa and Storey, 2021)

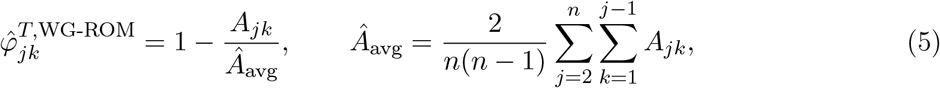

where *A*_*jk*_ is as in Eq. (1). Its biased limit is also a function of the true kinship matrix:

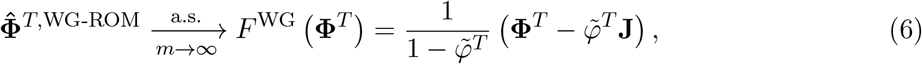

where 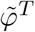 is the mean kinship excluding the matrix diagonal:

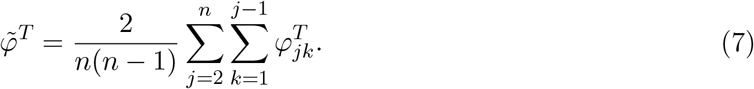

In Appendix C we prove that

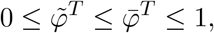

and equalities are achieved if and only if all kinship values are equal. Since the WG-ROM estimator closely resembles the popkin estimator in Eq. (1), it is clear that they are related by the bias function *F* ^WG^, while WG-MOR is introduced here and defined by the below formula:

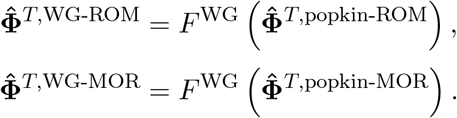

### 2.3 Association models

LMM and PCA are closely related association models (Astle and Balding, 2009; Hoffman, 2013; Yao and Ochoa, 2022):

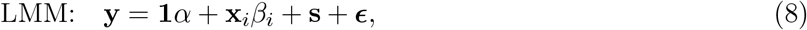

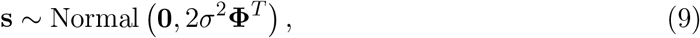

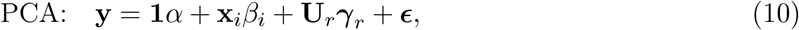

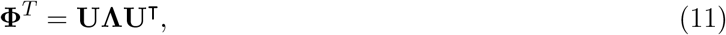

where **y** is a length-*n* vector of continuous trait values, *α* is the intercept coefficient, *β*_*i*_ is the genetic effect (association) coefficient of locus *i*, **s** is the (genetic) random effect, *σ*^2^ is the random effect variance factor, **U**_*r*_ is the *n* × *r* matrix of top-*r* eigenvectors (PCs) of **Φ**^*T*^, ***γ***_*r*_ is a length-*r* vector of coefficients for each eigenvector, 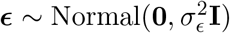 are random independent residuals, and **I** is the *n* × *n* identity matrix. Furthermore, Eq. (11) is the complete eigendecomposition of **Φ**^*T*^, where **U** is the *n* × *n* matrix of eigenvectors, and **Λ** is the *n* × *n* diagonal matrix of eigenvalues. As **s** and **U**_*r*_ play analogous roles in modeling the effect of relatedness in LMM and PCA, respectively, we refer to them jointly as relatedness effects, and *σ*^2^ and ***γ***_*r*_ as their coefficients.

### 2.4 Simulations

#### 2.4.1 Admixed family genotype simulation

An admixed family is simulated following previous work (Yao and Ochoa, 2022), except here only *K* = 3 ancestries are simulated and *F*_*ST*_ = 0.3 for the admixed individuals, which more closely resembles Hispanics and African Americans. Briefly, our admixture model first simulates *n* = 1000 founder individuals with *m* = 100, 000 loci. Random ancestral allele frequencies 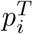, subpopulation allele frequencies 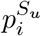, individual-specific allele frequencies *π*_*ij*_, and genotypes *x*_*ij*_ are drawn from this hierarchical model:

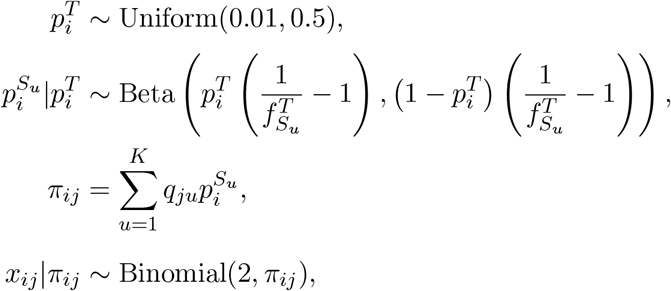

where this Beta is the Balding-Nichols distribution (Balding and Nichols, 1995) with mean 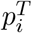 and variance 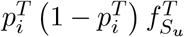. This is implemented in the R package `bnpsd`.

We also include family structure in the simulation. 20 generations are generated iteratively. Individuals in the first generation (*n* = 1000) are ordered by 1D geography, randomly assigned sex, and treated as locally unrelated. From the next generation, individuals are paired iteratively: randomly choosing males from the pool and pairing them with the nearest available female with local kinship < 1/4^3^ (to preserve the admixture structure) until there are no available males or females. Family sizes are drawn randomly ensuring every family has at least one child. Children are reordered by the average coordinates of their parents, their sex are assigned randomly, and their alleles are drawn from parents independently per locus. The simulation is implemented in the R package `simfam`.

#### 2.4.2 Trait simulation algorithm

Given an *m* × *n* genotype matrix 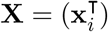, traits are simulated from

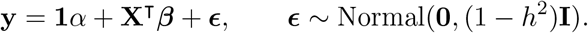

Given a desired number of causal loci *m*_1_ = *n/*10 and heritability *h*^2^ = 0.8, the goal is to choose causal coefficients ***β*** and the intercept *α* that result in zero mean and the desired trait heritability. Here, we use the “fixed effect sizes” trait simulation model described in (Yao and Ochoa, 2022). Briefly, first *m*_1_ causal loci are randomly selected, and for these steps only **X** is subset to these loci and reindexed. For known 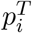, causal coefficients are constructed as:

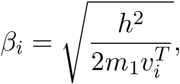

where 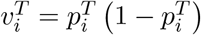; for unknown 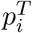 (real genotypes), the unbiased estimate 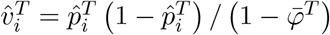 is used, where 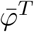 is the mean kinship estimated from `popkin`. Coefficients are made negative randomly with probability 0.5. For known 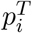, we obtain the desired zero trait mean with *α* = −2 (**p**^*T*^)^⊤^ ***β***, where here 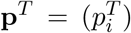 contains causal loci only. When 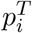 are unknown, to avoid covariance distortions, the intercept coefficient is constructed as

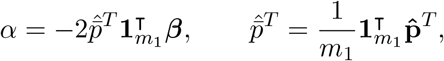

where 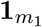 is a length-*m*_1_ column vector of ones.

### 2.5 Real genotype data processing

To evaluate different kinship estimators on a real dataset, we use the high-coverage NYGC version of the 1000 Genomes Project (Fairley et al., 2020), which is processed as before (Yao and Ochoa, 2022). Briefly, using `plink2` (Chang et al., 2015) we keep only autosomal biallelic SNP loci with filter “PASS”, pruned for linkage disequilibrium with parameters “`--indep-pairwise 1000kb 0.3`” to remove loci that have a greater than 0.3 squared correlation coefficient with other loci within 1000kb, and lastly remove loci with minor allele frequencies < 0.01. The resulting data have *m* = 1, 111, 266 loci and *n* = 2, 504 individuals. Traits are simulated for this dataset with *m*_1_ = *n/*10 = 250 causal loci.

### 2.6 Evaluation of performance

AUC_PR_ and SRMSD_*p*_ are used to evaluate approaches as before (Yao and Ochoa, 2022). Briefly, SRMSD_*p*_ (Signed Root Mean Square Deviation) measures the difference between the observed null p-value quantiles and the expected uniform quantiles:

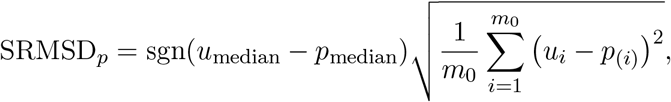

where *m*_0_ = *m* − *m*_1_ is the number of null (non-causal) loci, *i* indexes null loci only, *p*_(*i*)_ is the *i*th ordered null p-value, *u*_*i*_ = (*i*−0.5)*/m*_0_ is its expectation, *p*_median_ is the median observed null p-value, 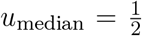 is its expectation, and sgn is the sign function (1 if *u*_median_ ≥ *p*_median_, -1 otherwise). SRMSD_*p*_ = 0 corresponds to calibrated p-values, SRMSD_*p*_ *>* 0 indicate anti-conservative p-values, and SRMSD_*p*_ < 0 are conservative p-values.

AUC_PR_ (Area Under the Precision and Recall Curve) is a binary classification measure that reflects calibrated power (Yao and Ochoa, 2022), which is calculated from the total numbers of true positives (TP), false positives (FP) and false negatives (FN) at some threshold or parameter *t*:

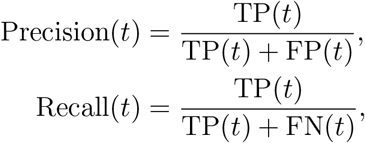

followed by calculating the area under the curve traced as *t* varies recall from zero to one. Higher AUC_PR_ is better, with best performance at AUC_PR_ = 1 for a perfect classifier, while worst performance at 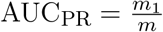 (overall proportion of causal loci) is for random classifiers.

### 2.7 Software

Popkin kinship estimates are calculated with the `popkin` R package. Standard MOR kinship estimates are calculated with GCTA (version 1.93.2beta). All other kinship estimators and limits are calculated using the `popkinsuppl` R package. PCs are calculated with the `eigen` function of R.

GCTA is used to run all LMM associations (Yang et al., 2011; Yang et al., 2014). We pass 2**Φ**^*T*^ for all kinship matrices tested (the same scale as its own kinship estimate). PCA association is performed with `plink2` (Chang et al., 2015). We use *r* = *K* − 1 = 2 PCs for the admixed family simulations, and *r* = 10 PCs for 1000 Genomes.

## 3 Results

### 3.1 Empirical analysis using admixed family simulation

To quantify the effect of kinship estimation bias, we simulate genotypes and traits, and calculate association p-values using a factorial design that tests all kinship matrix (3 bias types, times two locus weight types and one limit) and association model (PCA and LMM) combinations. We simulate an admixed population with *K* = 3 ancestries, who serve as founders for a 20-generation random pedigree. This high-dimensional admixed family scenario yields a large difference in performance between PCA and LMM (Yao and Ochoa, 2022).

Kinship estimates and limits for this simulation are shown in Fig. 1. The true kinship matrix shows the family relatedness as high values concentrated near the diagonal and the ancestry-driven population structure as the broad patterns off-diagonal. Only Popkin ROM is unbiased, while popkin MOR has a slight upward bias that varies across the matrix (Fig. S1A). In contrast, the Standard and Weir-Goudet (WG) estimates have large downward biases overall, resulting in abundant negative values; Standard biases vary for every pair of individuals, while WG has a uniform bias.

**Figure 1:**
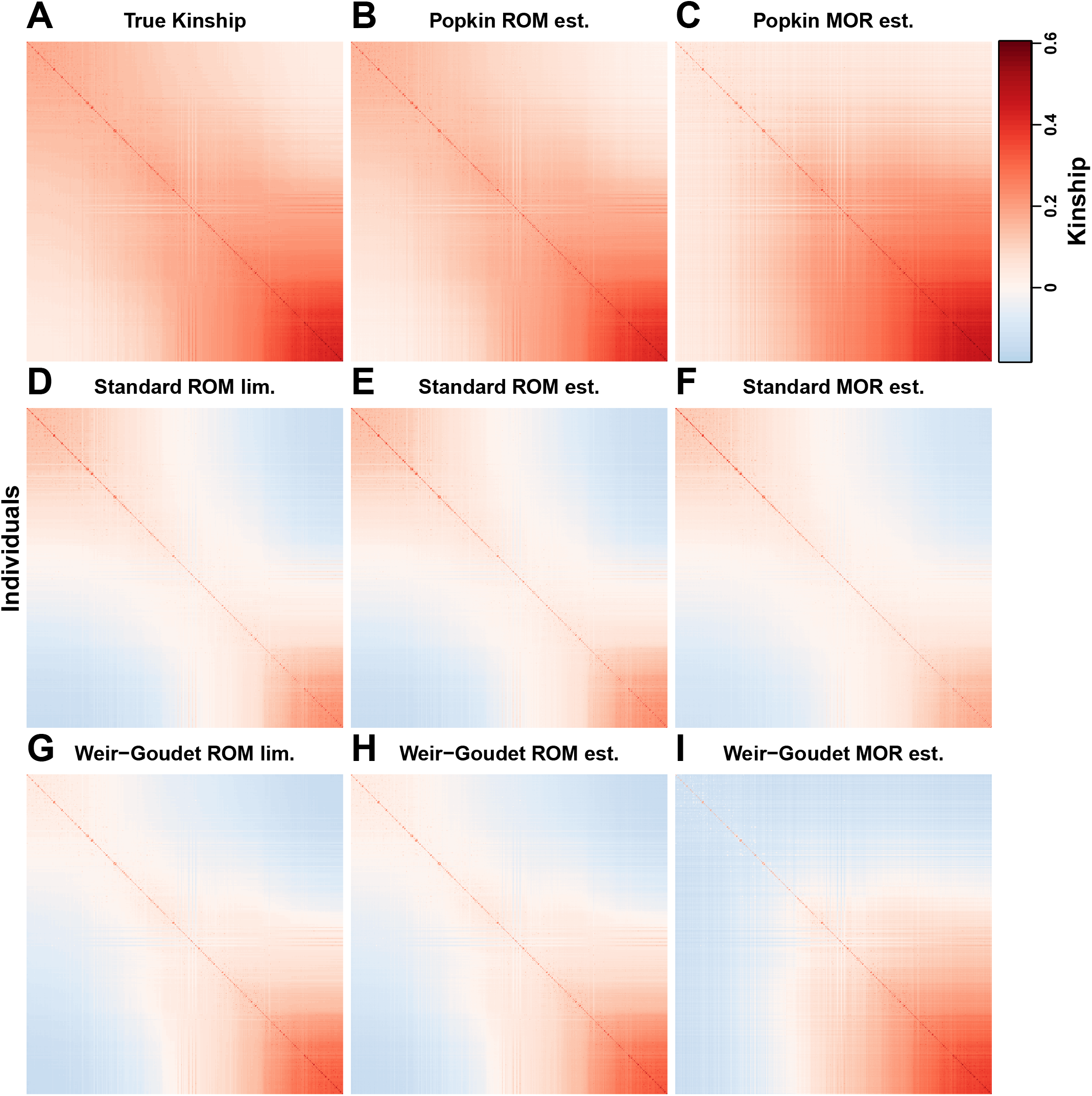
Kinship estimates and limits on the admixed family simulation. Each panel shows a kinship matrix as a heatmap, with each of the *n* = 1000 individuals along both x and y axes, color represents kinship: positive values in red, negative in blue. Diagonal contains inbreeding values. Each estimator bias type (Popkin, Standard, and Weir-Goudet; rows) has three matrices (columns): two locus weight types (ROM (ratio of means) and MOR (mean of ratios)) and limit of ROM.

We perform LMM and PCA association tests to determine how kinship biases affect association performance. Surprisingly, we find that kinship bias type does not have a discernible effect on association performance, as summarized by AUC_PR_ (a robust proxy for power; Fig. 2) and SRMSD_*p*_ (measures null statistic calibration; Fig. S2). The largest differences in performance are explained by the association model (LMM vs PCA), as expected due to our use of a family simulation where PCA performs poorly. Within association models, there are no clear differences between the performance of any of the kinship matrices, in fact many appear to have identical distributions (both statistics), the only clear exception being LMM popkin MOR, which has a few outlier replicates where performance is exceedingly poor.

**Figure 2:**
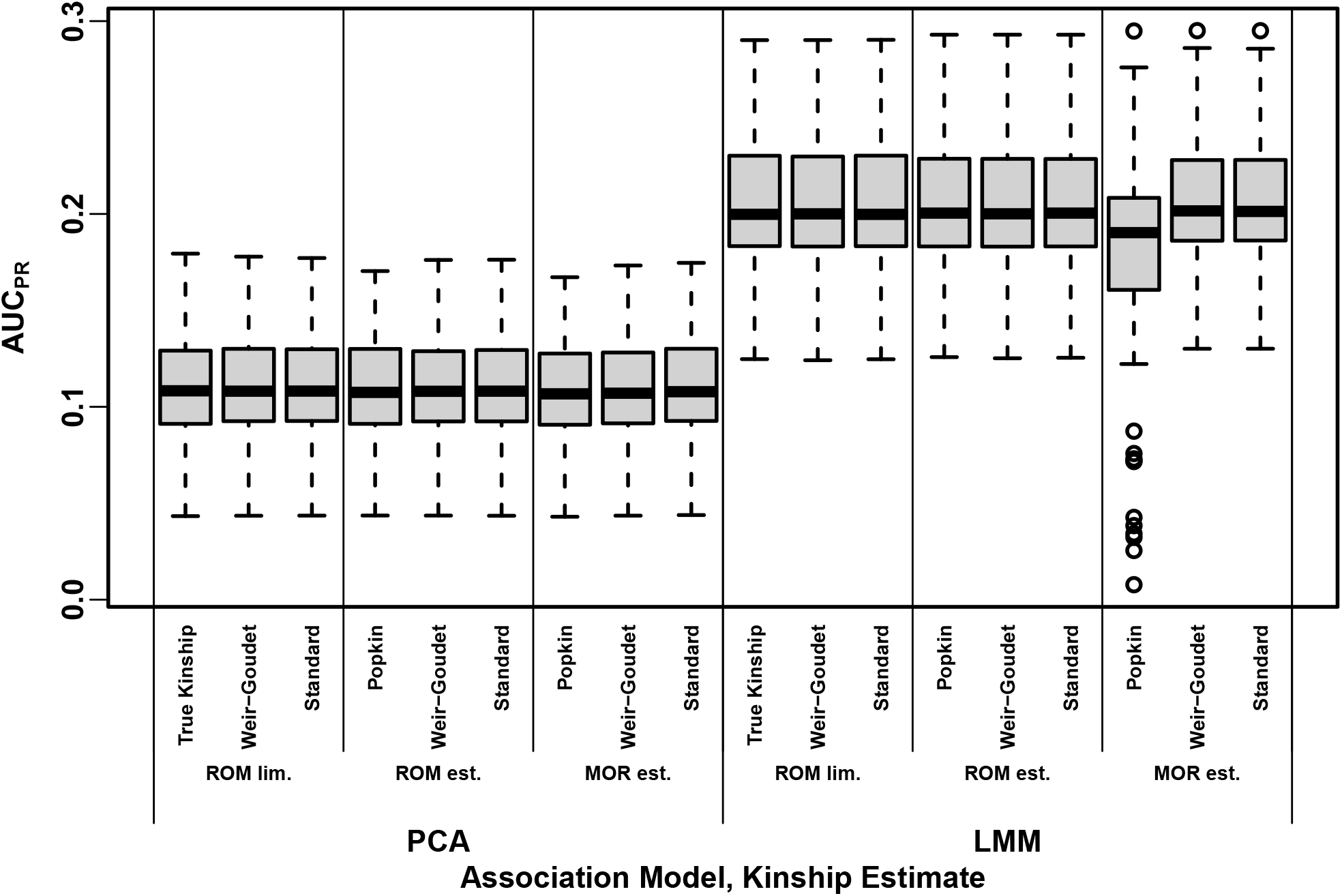
Distributions of Area Under the Precision-Recall Curve (AUC_PR_) on the admixed family simulation. Higher AUC_PR_ is better performance. Results for 100 replicates (each a random genotype matrix and trait vector). Approaches cluster primarily by association model (LMM or PCA), and vary little across bias types.

To better characterize the nearly identical performance distribution just observed, we next measure the agreement between individual association p-values. We calculate the proportion of loci between two methods with p-values within 0.01 of each other, which is an approximate measure of agreement, and find a remarkably high agreement between estimators of different bias types after matching association model and locus weight type or limit (Fig. 3). This is in contrast to low agreement between PCA and LMM statistics, and between LMM statistics with different locus weight types or limits. Minimum agreements are higher across PCA methods, though here the true kinship or popkin estimates disagree more from Standard and WG matrices. Overall, kinship matrices with different bias types (otherwise matched) result in nearly identical association statistics.

**Figure 3:**
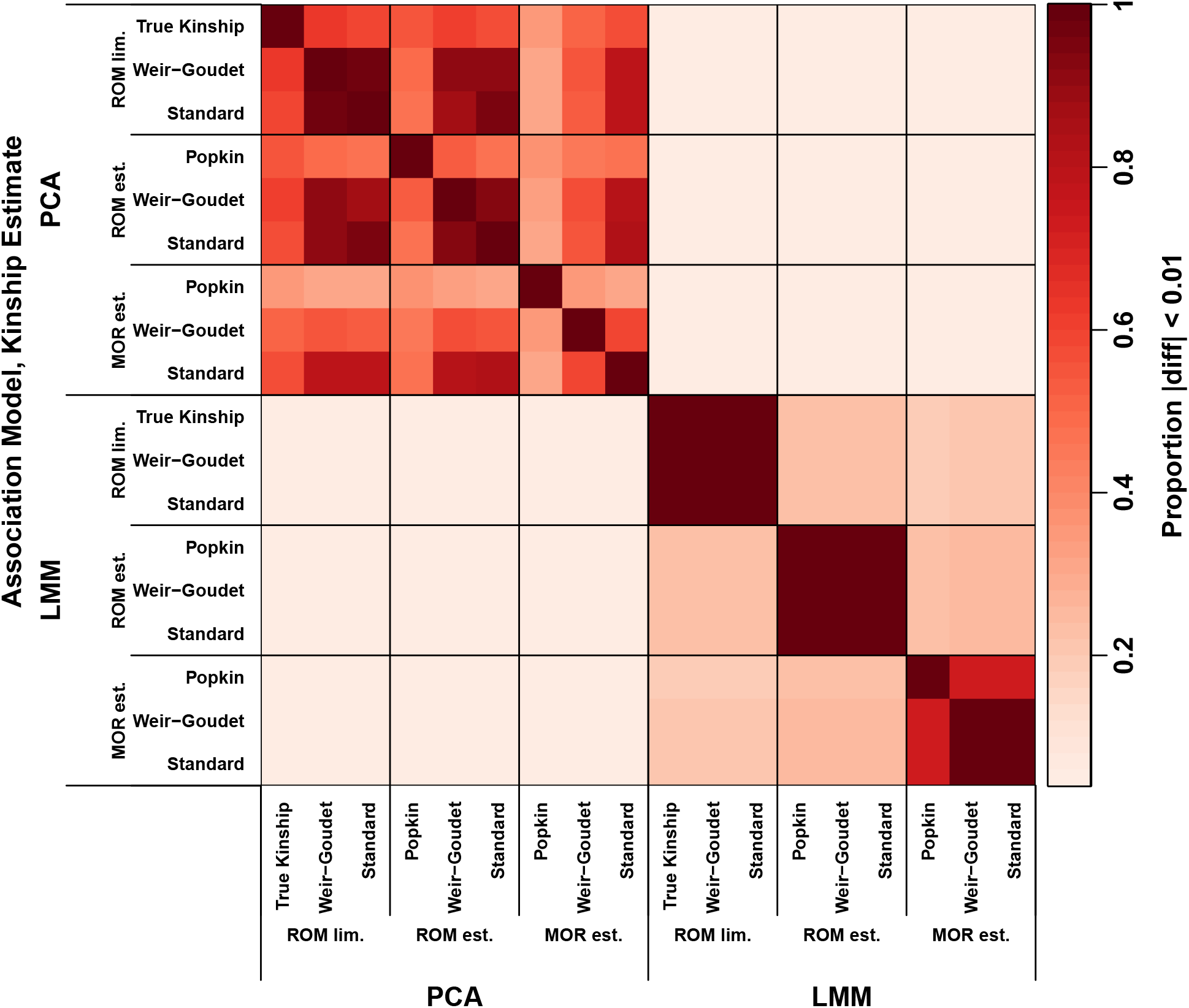
Approximate agreement between p-values on the admixed family simulation. Calculated agreement (absolute difference under 0.01) averaged over loci (color) of association p-values between association models (LMM vs PCA) and kinship matrices (x and y axes). All 100 replicates are used. Different bias types (matched for association model and locus weight type) have large proportions of nearly identical p-values.

### 3.2 Empirical analysis using 1000 Genomes

Now we repeat our analysis using the real genotypes of 1000 Genomes. Kinship estimates are shown in Fig. 4 (note real data have no true kinship or estimator limits). Popkin ROM estimates display an approximate nested block structure that arises from the tree relationships between subpopulations (Fig. 4A; trees were explicitly fit to this data in previous work (Yao and Ochoa, 2022)). However, popkin MOR estimates do not follow the nested blocks tree structure, since kinship between African and non-African populations is higher than kinship within African populations (Fig. 4B). Standard estimates have values are closer to zero, and a different bias for each pair of individuals, resulting in higher relative kinship for African compared to non-African populations (Fig. 4C-D). Lastly, WG estimates are uniformly smaller than popkin’s and attain large negative values (Fig. 4E-F).

**Figure 4:**
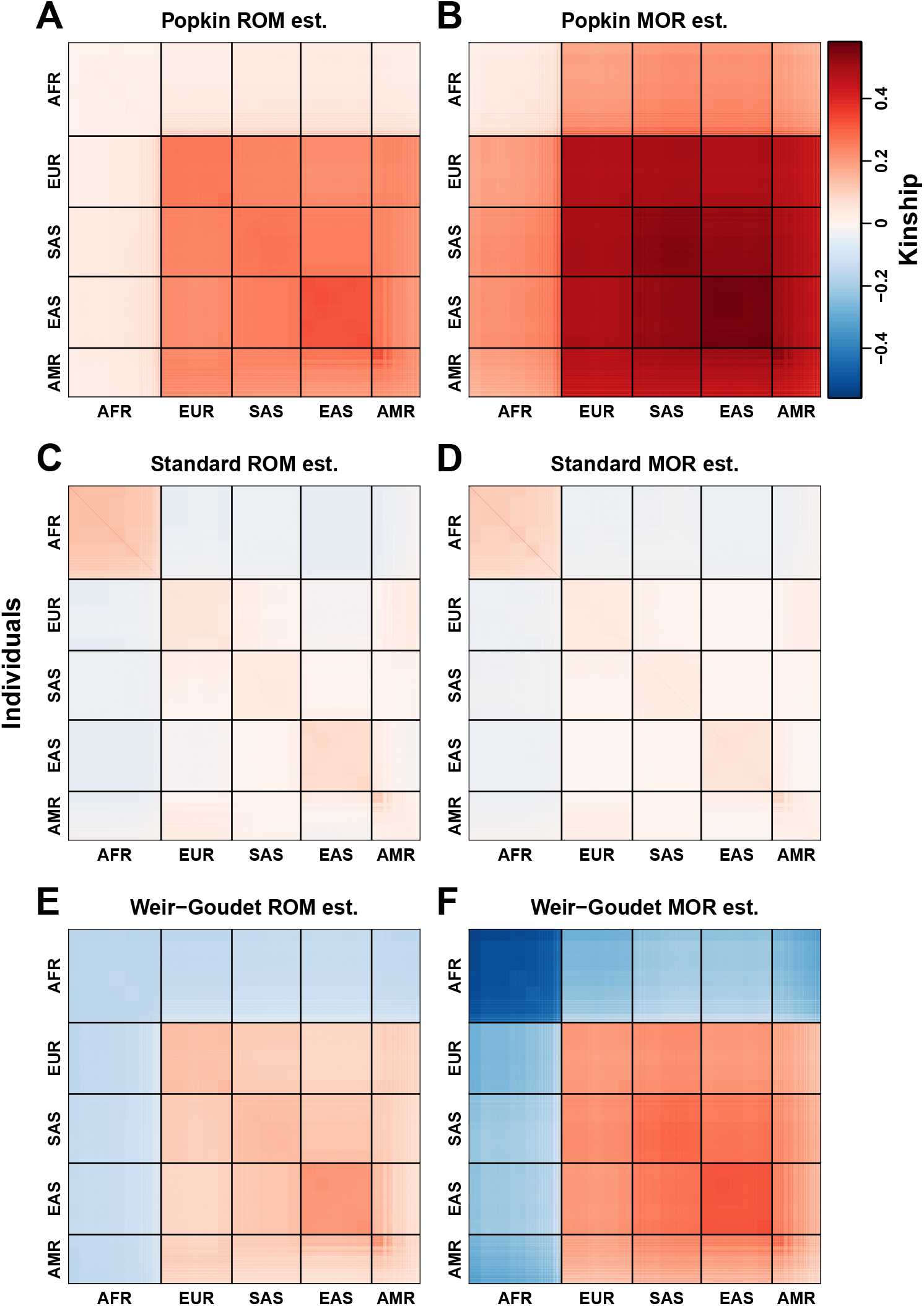
Kinship estimates on 1000 Genomes. Each panel represents a kinship matrix as a heatmap, as in Fig. 1. Superpopulation codes: AFR = African, EUR = European, SAS = South Asian, EAS = East Asian, AMR = Admixed Americans (Hispanics). Each estimator bias type (Popkin, Standard, and Weir-Goudet; rows) has two locus weight types (columns): ROM (ratio of means) and MOR (mean of ratios). In this visualization the upper range of all panels is capped to the 99 percentile of the diagonal (population inbreeding values) of the popkin MOR estimates.

Our association test conclusion are similar to our simulation study: AUC_PR_ and SRMSD_*p*_ distributions are nearly identical for estimators of different bias types but same locus weight type (ROM or MOR) and association model. However, unlike the simulation, here the MOR estimates noticeably outperform ROM estimates (LMM only), in terms of both AUC_PR_ (Fig. 5) and SRMSD_*p*_ (Fig. S3). P-values are again nearly identical at a large proportion of loci between approaches with matched association model and locus weight type (MOR or ROM), regardless of bias type (Fig. S4).

**Figure 5:**
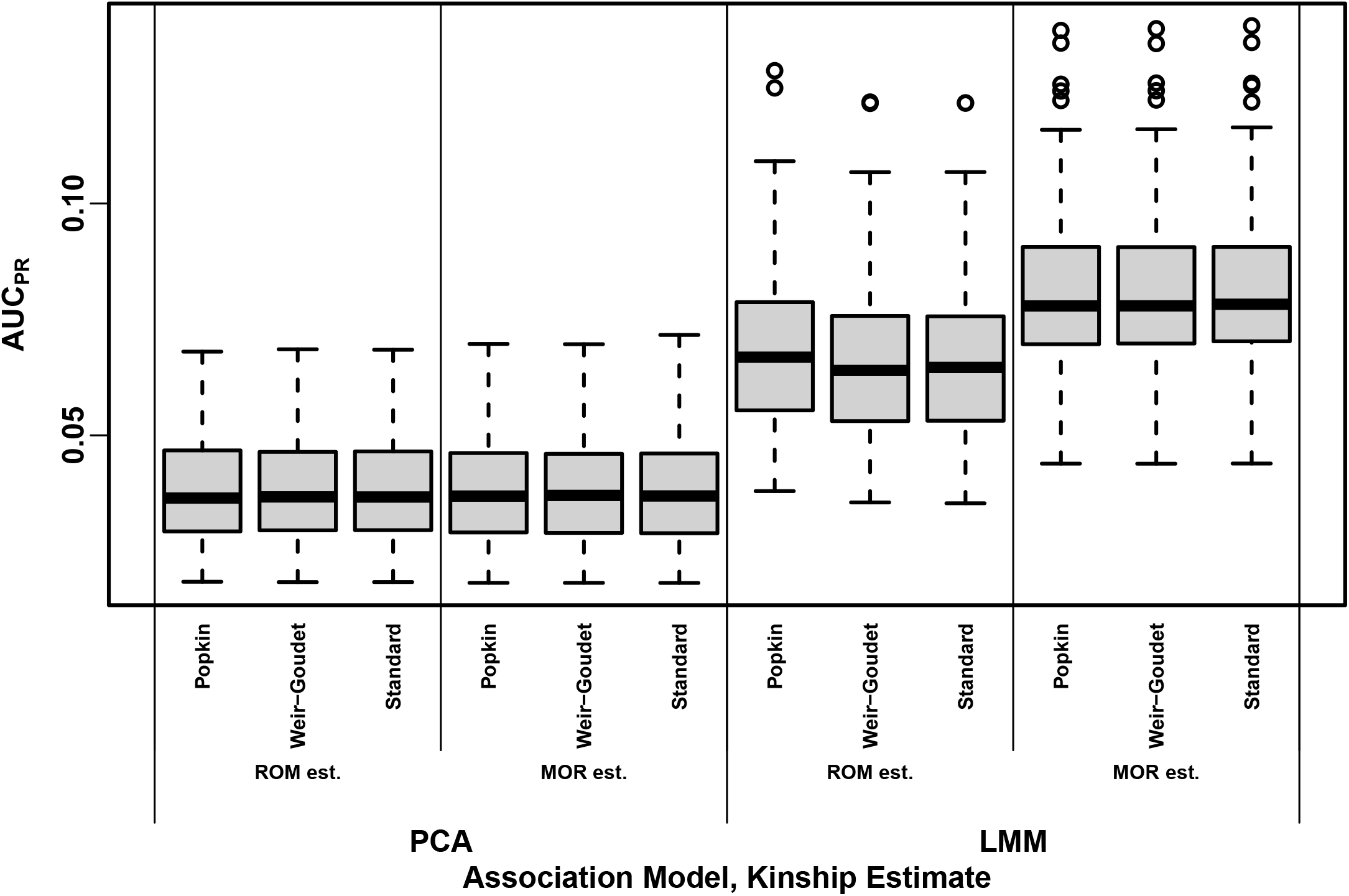
Distributions of Area Under the Precision-Recall Curve (AUC_PR_) on 1000 Genomes. Higher AUC_PR_ is better performance. Results based on 100 simulated trait replicates (real genotype matrix is fixed). Approaches cluster primarily by association model (LMM or PCA) and locus weight type (ROM or MOR), and do not depend much at all on the bias type.

### 3.3 Proof of association invariability to common kinship biases

Our empirical observations suggest that replacing a kinship matrix with either the Standard or WG-biased version does not alter association statistics (with exceptions we attribute to numerical precision artifacts); here prove a more general version of these facts mathematically. Our constructive proof shows that only a regression model with relatedness effects as covariates and an intercept is required, whose coefficients adapt to the bias, and no other coefficients change. This is fortunate, as the intercept and relatedness effect coefficients are nuisance parameters that usually go unreported, while the focal genetic association coefficient and its p-value are unchanged by these biases.

The most general form we identified of the bias function, mapping a kinship matrix to its bias-transformed version, and for which association invariability holds, is

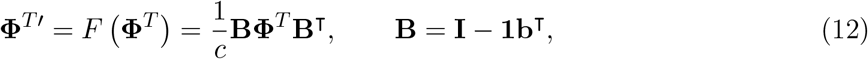

where *c* is any positive scalar and **b** is any length-*n* vector. The key property that the linear operator **B** must satisfy is that it shifts the input vector by the same scalar across its values, or

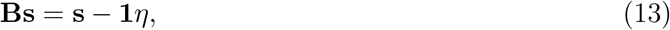

where **s** is any vector and the scalar *η* = **b**^⊤^**s** is a function of the input vector. **B** in Eq. (12) is the only form that results in Eq. (13).

The Standard bias function *F* = *F* ^std^ of Eq. (4) can be written as Eq. (12) with 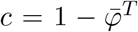 and 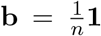 **1**, in which case **B** equals the centering matrix. Further, the generalized Standard estimator studied in Ochoa and Storey (2021) has **b** be a vector of individual weights that sum to one: **b**^⊤^**1** = 1. These **B** and **Φ**^*T*^′ are singular transformations (they are not invertible and have a zero eigenvalue), since **B1** = **0** and **B**^⊤^**b** = **0**.

The WG bias function *F* = *F* ^WG^ of Eq. (6) can be written as Eq. (12) with 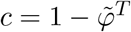 and

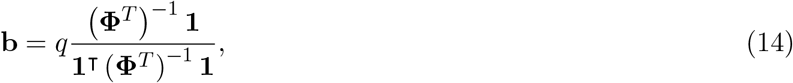

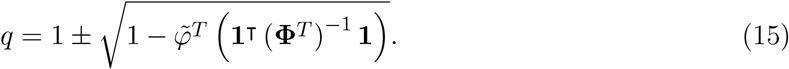

The derivation of this factorization is given in Appendix D. The determinant of the quadratic solution *q* would be non-negative if 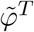 satisfied 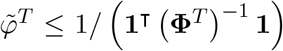. However, the actual 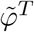 does not satisfy this inequality in any of our empirical cases, and in fact 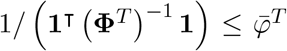 holds (proven in Appendix E; although 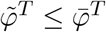 (Appendix C), in practice those two are very close while 1/(**1**^**⊤**^ (**Φ**^*T*^)^−1^ **1**) is much smaller than both), so *b* above is complex. This is a consequence of WG estimates being non-PSD, which we elaborate in the following sections. Nevertheless, PCA as well as the GCTA algorithms work for non-PSD matrices without invoking complex numbers (following sections and Appendix F).

#### 3.3.1 Proof for LMM case

Consider a random effect **s** drawn using **Φ**^*T*^, as given in Eq. (9). Using the affine transformation property of Multivariate Normal distributions (which holds even if **B** below is singular) and Eq. (12), then

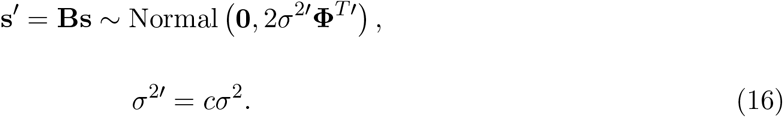

(This **s**′ has a degenerate distribution for Standard bias, since **Φ**^*T*^′ is singular, but **s**′+ ***ϵ*** is usually non-degenerate, since its covariance 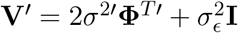 is invertible as long as 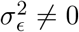.) Replacing **Bs** with the shift form in Eq. (13) shows that **s**′= **s** − **1***η* are equal in distribution. Therefore, the random effect **s**′of the biased kinship matrix differs from the random effect **s** of the original kinship only by **1***η*, a difference compensated for by adjusting the intercept coefficient in Eq. (8):

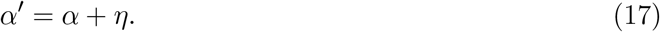

No other regression coefficients, or the total residuals, change when **Φ**^*T*^ is replaced with **Φ**^*T*^′, including the association coefficient *β*_*i*_ that is the focus of the test.

The above results require PSD kinship matrices **Φ**^*T*^′, which covariance matrices must be, and which are characterized by non-negative eigenvalues and determinants. Nevertheless, for the non-PSD WG bias (has a negative eigenvalue) combined with the generalized least squares association algorithm, which is used by GCTA and other LMMs (Kang et al., 2008; Kang et al., 2010; Yang et al., 2014), we find a stronger result consistent with Eq. (17), namely that *α*′= *α*, or in other words, *η* = 0 (Appendix F).

The LMM association p-value does not change in several common tests, including the F-test, since it only depends on the residuals and these do not change, as well as the likelihood ratio test, because although covariance determinants change, they cancel out in the ratio. The Wald test used by GCTA (Yang et al., 2014) is also invariant to these kinship biases given our empirical results in Figs. 3 and S4 and proven explicitly for WG bias in Appendix F. Lastly, we confirmed empirically that the Score test for the GCTA model is also invariant to these kinship biases (not shown). These arguments hold whether variance components are fit with maximum likelihood or restricted maximum likelihood (Kang et al., 2008; Kang et al., 2010; Yang et al., 2014), since multiplying the estimated genetic variance component *σ*^2^ by *c* and adjusting the intercept compensates for the bias regardless of how *σ*^2^, 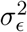 are estimated.

#### 3.3.2 Proof for PCA case

We present a proof for the PCA case that relies on an approximation that holds well in practice. Based on the PCA model of Eqs. (10) and (11), let **U**_*r*_ be the top eigenvectors of **Φ**^*T*^, and 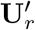 those of **Φ**^*T*^′. They key approximation is that

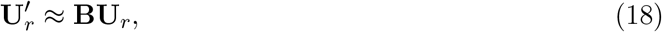

which is not strictly equal (since **BU**_*r*_ is not generally orthogonal, as eigenvectors must be), but we have found it to be a good approximation in practice. In this case the eigenvector coefficients need not change, 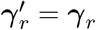, since the difference in scale of the kinship matrices (*c* in Eq. (12)) is absorbed by the eigenvalues not present in this model. Applying the shift of Eq. (13) shows that

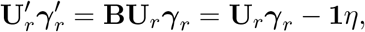

where 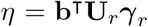 is a scalar. Therefore, the relatedness effects again differ only by **1***η*, which is compensated for by adjusting the intercept using Eq. (17), so the association coefficient *β*_*i*_ and the residuals are the same in both cases. This proof works if there are small numbers of zero or negative eigenvalues in **Φ**^*T*^′(non-PSD cases), as those rank last and are simply ignored. The observations from LMMs, that p-values are invariant to bias types, also hold for PCA.

We visualize the top PCs of our datasets in Fig. 6 to assess the validity of Eq. (18). The approximation is equivalent to each biased PC (Standard or Weir-Goudet) being shifted from the unbiased PC (Popkin), as described in Eq. (13). Fig. 6 indeed shows that PC1 is shifted by noticeable amounts in each of these cases, while PC2 is less shifted. However, a rotation of the PCs is also noticeable, particularly in the simulated data, and other large differences between MOR estimators, as expected since we know the approximation cannot be exact. Also, PCs can change order upon bias transformation, which we notice in the admixed family simulation, where PC2 and PC3 from popkin (and true kinship) actually correspond to PC1 and PC2, respectively, in both Standard and WG, and are plotted as such. No PC reordering occurs in 1000 Genomes. Overall, while the approximation of Eq. (18) can be weakened to merely require that the biased PCs plus intercept span the same subspace of the unbiased PCs plus intercept, the approximate PC shifts better explain intuitively why the result for LMM is also observed for PCA association.

**Figure 6:**
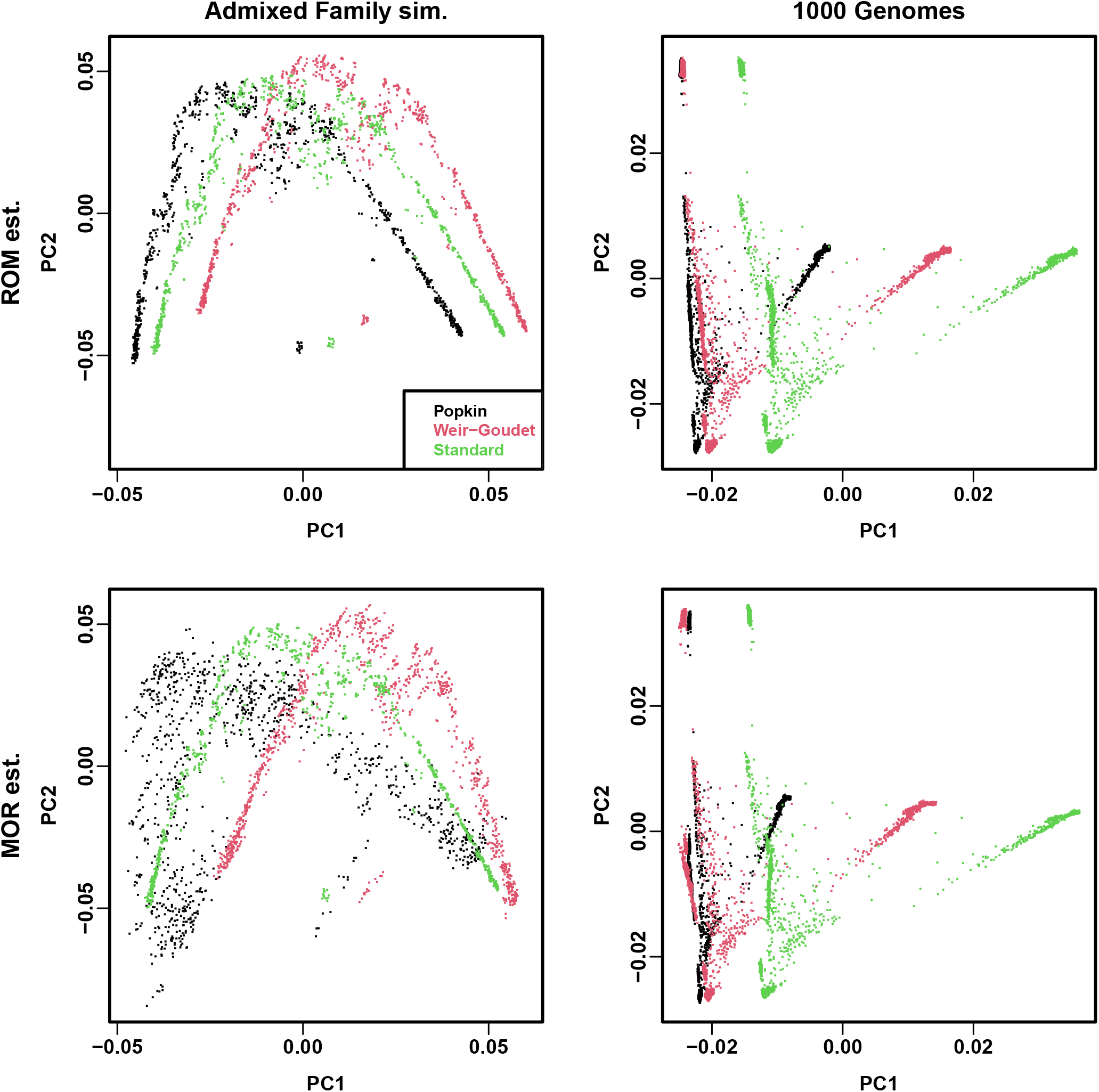
Visualization of PC shift due to kinship biases. Each panel shows three estimates (bias types): Popkin, Standard, and Weir-Goudet. ROM estimates are in first row, MOR in second row. (In admixed family, ROM limits are very similar to ROM estimates (not shown).) Columns show estimates from each dataset: admixed family simulation (first replicate) and 1000 Genomes. For popkin (both ROM and MOR estimates) in admixed family only, PC1 and PC2 are replaced with PC2 and PC3 (see text).

### 3.4 Proof of association invariability to change in ancestral population

The kinship matrices we used so far have values that depend on the choice of ancestral population *T*. Here we consider the effect on association of changing ancestral population, and prove that it is also compensated for by the relatedness and intercept coefficients.

Start from a kinship matrix **Φ**^*S*^ in terms of ancestral population *S*, and let *T* is a population ancestral to *S*. If the inbreeding coefficient of *S* when *T* is the reference ancestral population is 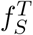, then the kinship matrix **Φ**^*T*^ in terms of *T* is given by (Ochoa and Storey, 2021)

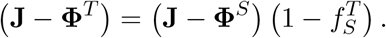

Solving for **Φ**^*T*^ and simplifying results in

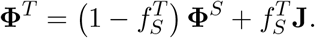

This resembles WG bias but in reverse: whereas WG reduces and rescales kinship by 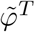, changing to a more ancestral population rescales and increases kinship by 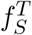. Indeed, excluding 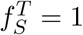, this transformation can be written as Eq. (12) with 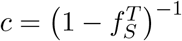 and

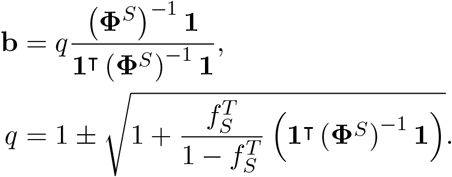

The determinant of *q* is strictly positive, since **1**^T^ (**Φ**^*S*^)^*−*1^ **1** *>* 0 (since **Φ**^*S*^ is positive definite, its inverse is too) and 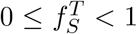. Thus, our previous results apply: ancestor change is compensated for by the relatedness and intercept coefficients, so the association statistics are invariant to this transformation.

### 3.5 Characterization of non-PSD and singular kinship and trait covariance estimators

While attempting to validate and characterize the earlier factorization of the WG bias function (Eqs. (12) to (15)), we discovered that it does not produce PSD matrices, which covariance matrices are required to be. To characterize this problem more broadly, we calculate the eigenvalues of all kinship matrices **Φ**^*T*^ and trait covariance matrices 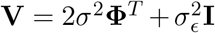, the latter used by LMMs and which we calculate using GCTA’s estimates of *σ*^2^ and 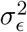.

We find that all WG matrices have very large negative minimum eigenvalues, and popkin MOR estimates also have smaller negative minimum eigenvalues (Fig. S5). Moreover, besides all WG matrices and most popkin MOR estimates, Standard matrices are also often non-PSD but only in 1000 Genomes (Fig. 7), which has missing genotypes (the admixed family simulation does not have missing genotypes). Each of these non-PSD matrices only has one negative eigenvalue. Notably, all popkin ROM estimates are PSD in every evaluation, including under missingness in 1000 Genomes.

**Figure 7:**
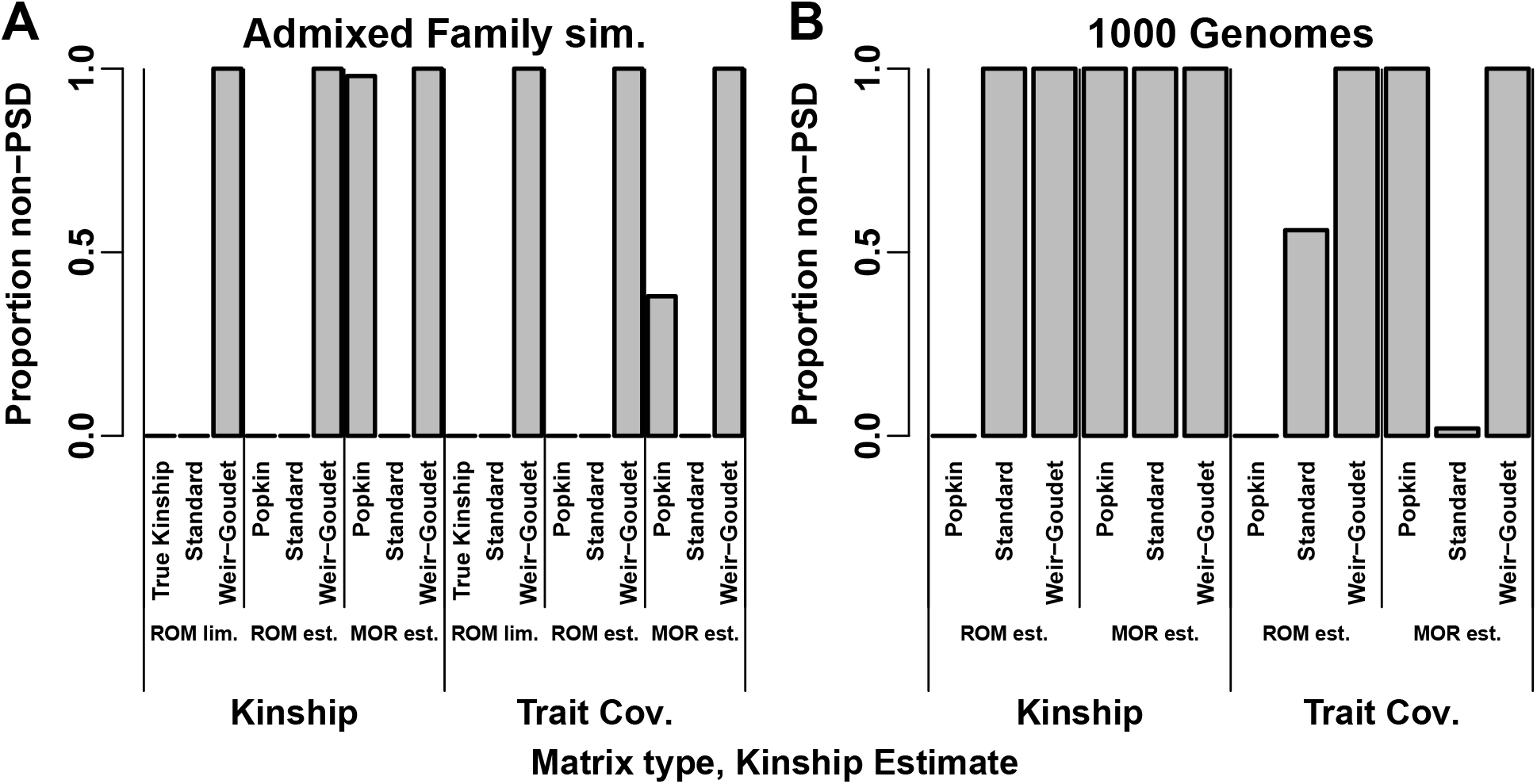
Proportion of kinship and trait covariance (V) matrices that are not positive semidefinite (PSD). A matrix is non-PSD if it has negative eigenvalues (below − 10^*−*7^ to allow for limited machine precision). Proportion is calculated over 100 replicates (1000 Genomes kinship has one value since genotypes are fixed, but **V** varies per replicate). **A**. In admixed family simulation, which does not have missing genotypes, all WG matrices and most popkin MOR estimates are non-PSD. All non-PSD kinship matrices results in non-PSD **V** except some popkin ROM estimates yield PSD **V. B**. In 1000 Genomes, which has missingness, all kinship estimates are non-PSD except popkin ROM. Of the non-PSD kinship matrices, only some Standard estimates yield PSD **V**.

In order to quantify matrix singularity, as well as numerical accuracy problems caused by multiplying by inverses of nearly-singular matrices, we calculate condition numbers, which equal the maximum absolute eigenvalue divided by the minimum absolute eigenvalue of our covariance matrices. As expected, we see that Standard kinship matrices are singular on our admixed family simulation (which lacks missingness), as reflected by extremely high condition numbers, but their trait covariances have small condition numbers (Fig. S6). No other matrices are singular, but popkin MOR estimates in the admixed family simulation have relatively high condition numbers for both kinship and trait covariance.

Consider the theoretical connection between the eigenvalues of **Φ**^*T*^ and those of **V**. The eigendecomposition trick widely used to fit variance components in LMMs (Kang et al., 2008; Lippert et al., 2011; Svishcheva et al., 2012; Zhou and Stephens, 2012; Sul et al., 2018) yields

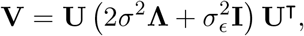

where **U** and **Λ** are the eigenvectors and eigenvalues of **Φ**^*T*^, respectively (Eq. (11)), so the eigenvectors of **V** are also **U** and its eigenvalues are 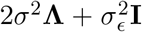. Therefore, since 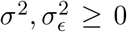, then if **Φ**^*T*^ is positive definite (all of its eigenvalues are positive) then so is **V**, and the condition number of **V** is always smaller (better) or equal than that of **Φ**^*T*^. A negative kinship eigenvalue *λ*_*k*_ may become positive for **V** only if 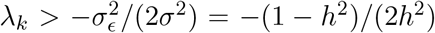, so very large negative *λ*_*k*_ values as observed for WG do not become positive in **V**, in fact they can become more negative (Fig. S5). **V** is always invertible and well-conditioned even when **Φ**^*T*^ is singular PSD (has zero eigenvalues), as the Standard estimator is under no missingness, since a kinship zero eigenvalue becomes 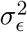 for **V**. Conversely, the above equation explains why some non-PSD kinship matrices are particularly problematic: negative eigenvectors near 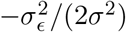 can result in ill-conditioned **V**. We see that popkin MOR estimates are non-PSD (Fig. S5) in such a way that some of their **V** are ill-conditioned (Fig. S6), and this explains its poorer performance in the admixed family evaluations (Figs. 2 and S2), as shown in the next subsection.

### 3.6 Further empirical validation of theoretical predictions

Seeing that WG is always non-PSD, and to query other instances where predictions are not fully met, here we analyze estimation accuracy of various parameters to better understand theoretically and empirically how broken assumptions affect them. With PCA, no deviations from expectation of AUC_PR_ and SRMSD_*p*_ are observed for WG (Figs. 2 and 5, Figs. S2 and S3), which makes sense since PCA simply ignores eigenvectors with negative or zero eigenvalues. Therefore, our analysis focuses on LMM, where deviations are observed and clarification regarding WG is needed.

LMMs such as GCTA perform association testing in two steps. First is the restricted maximum likelihood step used to fit variance components. Although the eigendecomposition approaches (Kang et al., 2008; Lippert et al., 2011; Svishcheva et al., 2012; Zhou and Stephens, 2012; Sul et al., 2018) require positive definite **V** (lest the determinant of **V** be negative), surprisingly the GCTA average information algorithm only requires in practice that **V** be invertible (Yang et al., 2011). Thus, the relationship between WG, Standard, and True or Popkin variance components are largely as expected from our theoretical prediction *σ*^2′^ = *cσ*^2^ in Eq. (16), with the exception of popkin ROM on 1000 Genomes only, whose genetic variance estimates are slightly smaller than expected (Fig. S7). Next we determine the effect of WG bias on coefficient estimates. In this second step of LMM association testing, once **V** is determined, GCTA and other LMMs use generalized least squares to estimate fixed effects coefficients (Kang et al., 2008; Kang et al., 2010; Yang et al., 2014). Using the first replicate of the admixed family simulation and the true kinship matrix and the Standard and WG limits only, we recalculate the genetic effect *β*_*i*_ and intercept coefficients *α* in R for all loci, and confirm that we recover the GCTA estimates for *β*_*i*_ to the given precision. We then compare intercept coefficients, which are not given by GCTA, and confirm our theoretical prediction (Appendix F) that they are identical whether the True or WG ROM limit kinship matrices are used (the mean absolute difference is below 10^*−*7^). In contrast, intercepts fit using the Standard ROM limit kinship matrix are different than those of the true kinship (not shown), which agrees with our theoretical prediction that the intercept varies to compensate for the kinship matrix bias (*α*′= *α*+ *η* in Eq. (17)).

Lastly, we explain the largest deviations from our predictions of the performance metrics AUC_PR_ and SRMSD_*p*_. We find that the small performance errors of popkin ROM in 1000 Genomes (Figs. S3 and 5) are driven by errors in genetic variance component estimation *σ*^2^ (Fig. S8). However, the larger performance errors of popkin MOR in the admixed family simulation (Figs. 2 and S2) are instead explained by the condition number of **V** (Fig. S9). This result makes sense since the condition number by definition quantifies regression coefficient estimation accuracy.

## 4 Discussion

Previous research showed that commonly used kinship estimators are biased, and that these biases can be large (Ochoa and Storey (2021); Fig. 1). Our initial hypothesis was that these kinship biases would affect association testing, but surprisingly found that association is unaffected. We then proved theoretically that it is the intercept and relatedness effect (random effect or PCs) coefficients that compensate for the bias, and result in identical association coefficients and significance statistics.

Kinship estimates depend on the choice of ancestral population, which conditions the distributions of allele frequencies and genotypes, but the effect of this choice of association testing was not only unknown but completely disregarded. A corollary of our theoretical results is that changes of ancestral population, which behave algebraically like kinship bias, are also compensated for by the relatedness and intercept coefficients, so association testing is also invariant to the choice of ancestral population. Thus, although a choice of ancestral population is always being made when estimating kinship, this choice is fortunately inconsequential to association testing, as it ought to be since relatedness is being conditioned upon in these tests.

Given that kinship bias type is not important for association studies, we are free to choose a kinship estimator based on other properties. Ideally, kinship matrices result in well conditioned trait covariance matrices, since that has the largest effect in numerical accuracy and power in LMMs. Well-conditioned association is guaranteed for PSD kinship matrices, and popkin ROM is the only estimator that produces PSD matrices consistently across our evaluations (Fig. 7). Popkin ROM is also the only unbiased kinship estimator (Ochoa and Storey, 2021). We observed that Standard kinship estimates are also not PSD when genotypes are missing, a well understood phenomenon for related sample covariance estimators outside genetics (Jurczak and Rohde, 2017). Fortunately, non-PSD kinship estimators often perform well for association. Nevertheless, in our admixed family simulation we did see the other popkin estimator (the MOR version) perform particularly poorly due to being non-PSD, which in combination with the heritability parameters of this simulation results in ill-conditioned association tests and substantial loss of accuracy and power (Figs. 2, S2 and S9). Theory predicts that the same can happen with any non-PSD estimator, depending on unknowns such as the heritability and the value of the negative eigenvalues of the kinship estimator, so it is risky to use MOR estimators (all of which are non-PSD in 1000 Genomes), as well as the WG estimator generally (which is non-PSD in all replicates of all of our evaluations). We also observe smaller numerical inaccuracies for popkin ROM, the estimator we recommend, in 1000 Genomes only, although the result is mixed: performance is slightly better (Fig. 5) although null p-value calibration is slightly worse (Fig. S3). The cause is variance components are poorly estimated (Fig. S7), but we did not find a more fundamental explanation. Overall, our assessment suggests that the popkin ROM estimator is the safest choice due to its guarantee of well-conditioned associations that other estimators cannot make.

Despite being non-PSD, we observe better performance for MOR versus ROM estimators in LMM association of 1000 Genomes (Fig. 5). Perhaps this is expected because we simulated larger coefficients for rare variants, while MOR estimators upweigh rare variants. This effect is not observed in the admixed family simulation, where MOR and ROM versions give similar kinship estimates (Fig. 1) and performed similarly (Fig. 2), compared to 1000 Genomes where kinship estimates are also strikingly different (Fig. 4). However, only popkin ROM is unbiased (Fig. 1B, Fig. S1). One potential explanation is that our kinship model assumes that all variants existed in the MRCA population, whereas rare variants in human data are known to be more recent mutations, and thus their effective kinship matrix is different than that of ancestral variants. Therefore, despite its biases, the popkin MOR estimator may better capture the covariance of rare variants and thus model them better in association tests, particularly in LMMs where the effect is most pronounced. Future work should focus on better approaches for upweighing rare variants or otherwise estimating their covariance structure while resulting in positive definite kinship estimates.

Our conclusions that common kinship biases do not affect association studies extend to variations of the Standard kinship estimator that weigh loci according to linkage disequilibrium (Speed et al., 2017; Wang et al., 2017), which also have the Standard bias type since this bias is present in each locus (Ochoa and Storey, 2021). As shown in our theoretical results, another form of the Standard kinship estimator that weighs individuals to estimate ancestral allele frequencies 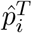, including the best unbiased linear estimator in Appendix E (Astle and Balding, 2009; Thornton and McPeek, 2010), is also subject to the same conclusions.

In this study, we show empirically and theoretically that association tests are invariant to the use of common kinship estimators that are biased versus a more recent unbiased estimator. The underpinnings of our proof show that the same result holds for association with generalized linear models, since the intercept and relatedness effects interact in the same way as for linear models (the link function goes around the trait only); these models include case/control models such as logistic PCA and LMM. However, heritability estimation requires unbiased estimates of the random effect coefficient (*σ*^2^), so it is biased when the standard kinship estimator is used, as it is using GCTA (Yang et al., 2011; Yang et al., 2014). Nevertheless, heritability estimation is a complex problem and a complete analysis is beyond the scope of this work. Overall, we have described an unexpected robustness of association studies, and our theoretical understanding of this result may help guide future improvements for association and other related models.

## Abbreviations

PCA: principal component analysis
PCs: principal components
LMM: linear mixed-effects model
MOR: mean of ratios
ROM: ratio of means
WG: Weir-Goudet (kinship estimator)
MRCA: Most Recent Common Ancestor
SRMSD_*p*_: p-value Signed Root Mean Square Deviation
AUC_PR_: Area Under the Precision Recall Curve
GCTA: Genome-wide Complex Trait Analysis (software)
PSD: positive semidefinite

## Declaration of interests

The authors declare no competing interests.

## Acknowledgments

This work was funded in part by the Duke University School of Medicine Whitehead Scholars Program, a gift from the Whitehead Charitable Foundation. The 1000 Genomes data were generated at the New York Genome Center with funds provided by NHGRI Grant 3UM1HG008901-03S1.

## Web resources

plink2, https://www.cog-genomics.org/plink/2.0/

GCTA, https://yanglab.westlake.edu.cn/software/gcta/

bnpsd, https://cran.r-project.org/package=bnpsd

simfam, https://cran.r-project.org/package=simfam

simtrait, https://cran.r-project.org/package=simtrait

popkin, https://cran.r-project.org/package=popkin

popkinsuppl, https://github.com/OchoaLab/popkinsuppl

## Data and code availability

The data and code generated during this study are available on GitHub at https://github.com/OchoaLab/bias-assoc-paper. The high-coverage version of the 1000 Genomes Project was downloaded from ftp://ftp.1000genomes.ebi.ac.uk/vol1/ftp/data_collections/1000G_2504_high_coverage/working/20190425_NYGC_GATK/.

## Appendices

### A Justification for popkin generalizations

The popkin estimator in Eq. (1) has been generalized in this work to include locus weights *w*_*i*_. The original ROM formulation had *w*_*i*_ = 1 for all loci *i* (Ochoa and Storey, 2021). Recalling from that original work that

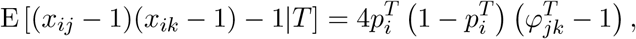

then for fixed *w*_*i*_ we get

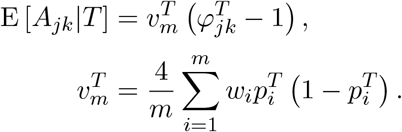

Therefore, as before all the unknowns 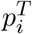 and now also the known weights *w*_*i*_ collapse into a single parameter 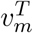, which is estimated under the assumption that the minimum kinship is zero, giving 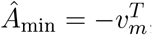, so that

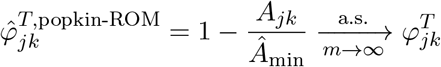

as desired.

The MOR case of 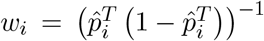 does not fit the previous case because this *w*_*i*_ is a random variable (it is a function of the genotypes). The term of interest *w*_*i*_((*x*_*ij*_ − 1)(*x*_*ik*_ − 1) − 1) is a ratio of random variables whose expectation does not have a closed form. In this case, we rely on the first-order approximation to this expectation, namely

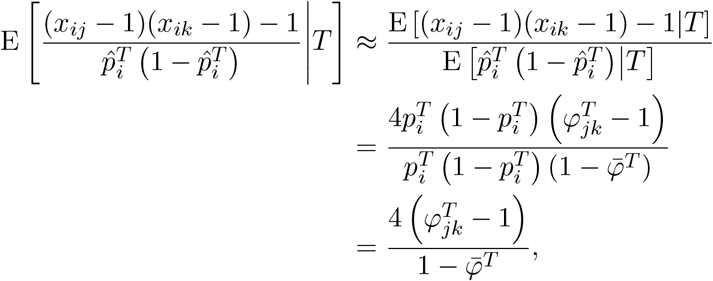

where the expectation of 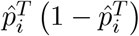 was calculated previously (Ochoa and Storey, 2021). In this case the expectation of *A*_*jk*_, summing across loci, is also approximated by

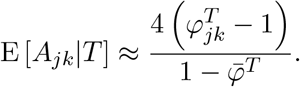

The same strategy as before applies to estimate the unknown factor 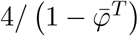, namely that if the minimum kinship is zero then 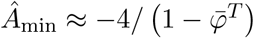, resulting in

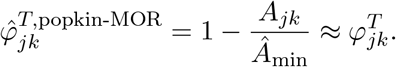

### B Connection between popkin and standard kinship estimator

Since the connection we discovered holds when data are complete, but not under missingness, to determine necessary conditions we introduce more complete forms of the estimators that handle missingness. Popkin (with locus weights) has the following parts updated:

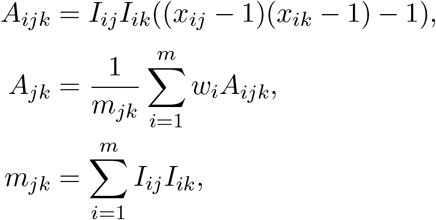

where *I*_*ij*_ = 1 if *x*_*ij*_ is not missing, 0 otherwise (this way missing *x*_*ij*_ can have any value and not contribute to the estimator). Only loci with both genotypes (*x*_*ij*_ and *x*_*ik*_) non-missing are included in the above average, and *m*_*jk*_ counts the total number of such loci. The ancestral allele frequency estimator with missingness is

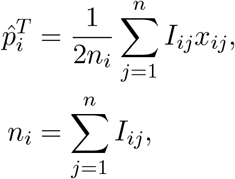

which averages over individuals rather than loci, so its denominator is the number of non-missing individuals at this locus. Let us compute some averages of the popkin estimator. Since the result we want holds at every locus separately, let us formulate the averages of interest at locus *i* only:

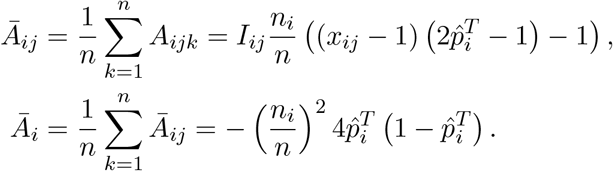

Therefore, the combination of interest is:

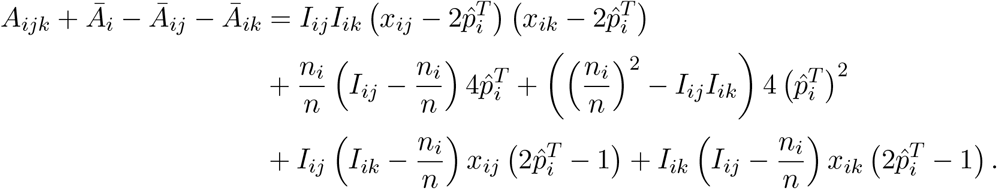

For the above to equal 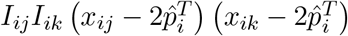, which is the first term above, the rest of the terms must vanish for arbitrary values of 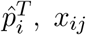, and *x*_*ik*_. Since *n*_*i*_ *>* 0 (there is at least one non-missing individual at every locus), the term 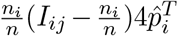 vanishes if and only if 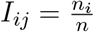, and since *I*_*jk*_ = 0 does not solve this equation (because *n*_*i*_ *>* 0), then *I*_*jk*_ = 1, which requires *n*_*i*_ = *n*, so no individuals can have missing data at this locus (the rest of the terms vanish when this is so).

Thus,

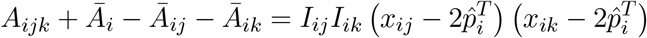

if and only if there is no missing data at locus *i*. The other desired result of

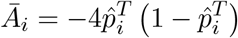

also requires *n*_*i*_ = *n*.

Assuming now no missingness, transforming the popkin estimates using the Standard bias function of Eq. (4) gives

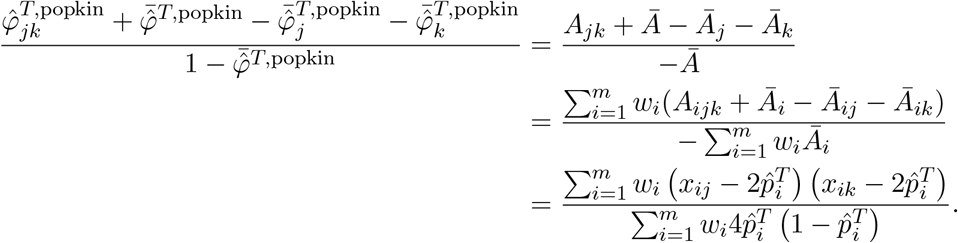

Therefore, if popkin ROM is input (*w*_*i*_ = 1), this transformation yields Standard ROM. On the other hand, if popkin MOR is used 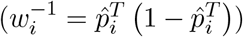, the transformation yields Standard MOR.

### C Mean kinship inequalities

Denote the mean of the diagonal kinship terms as 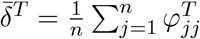. Here we prove that

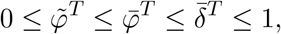

with each of 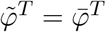 and 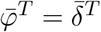 if and only if all kinship values are equal.

The inequalities 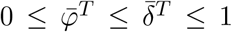 follow directly from previous work, applied to a kinship matrix rather than a coancestry matrix as done originally, as the proof required solely a covariance matrix with values between 0 and 1 (Ochoa and Storey, 2021). 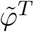 is defined in Eq. (7). 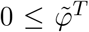 follows since every kinship value is non-negative. 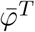 and 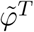 are related by

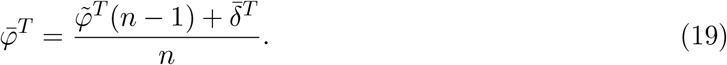

Applying 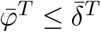 to Eq. (19) and simplifying yields 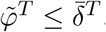. Lastly, since 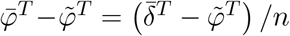 (from rearranging Eq. (19)), it also follows that 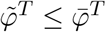, as desired. Furthermore, 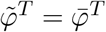 holds if and only if all 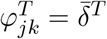, since that is necessary and sufficient for 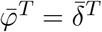.

### D Derivation of WG bias factorization

Here we rewrite the WG bias function of Eq. (6) as a factorization of the form of Eq. (12). It is easy to see that 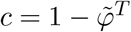. Expanding Eq. (12) gives

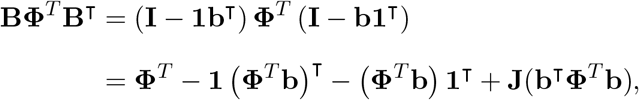

where **b**^⊤^**Φ**^*T*^ **b** is a scalar and **Φ**^*T*^ **b** a vector. Equating the above to Eq. (6) and rearranging, we obtain

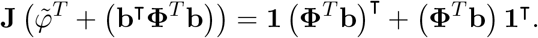

Since 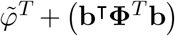 is a scalar and **J** = **11**^⊤^, we can see that the solution requires the right side to also be a constant matrix, which is only achieved if **Φ**^*T*^ **b** ∝ **1**. We choose the scaling factor for the last **1** to be 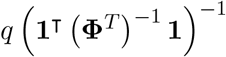 as this simplifies notation later, and solving for **b** results in Eq. (14). To solve for q, we replace **b** from Eq. (14) into the above equation, which after rearranging results in

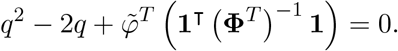

The solution to the above quadratic equation is given by Eq. (15), as desired.

### E Minimum weighted mean kinship

Consider the weighted mean kinship value **w**^⊤^**Φ**^*T*^ **w**, where **w** are weights that sum to one (**w**^⊤^**1** = 1). The ordinary mean kinship 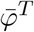 is the special case with 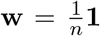. The weights that minimize the weighted mean kinship are the solution of the Lagrangian multiplier problem

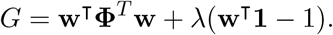

The derivatives are the constraint and 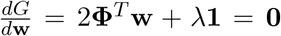. The optimal weights thus satisfy 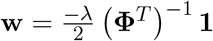. Multiplying by **1**^T^, since **1**^⊤^**w** = 1, allows us to solve for 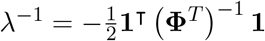. Thus, the optimal weights are

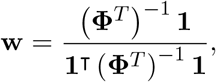

a solution that recurs in related settings, and applied to genotypes as 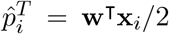 yields the best linear unbiased estimator of 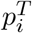 (Altschul et al., 1989; Astle and Balding, 2009; Thornton and McPeek, 2010). Therefore, the minimum weighted mean kinship is, and satisfies,

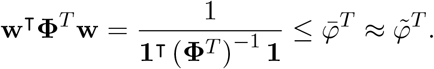

### F Proof that WG bias results in zero intercept shift under LMM generalized least squares estimation

For this section suppose that variance components have been estimated, so 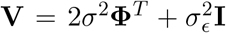 is given, assume it is invertible, and rewrite the LMM as

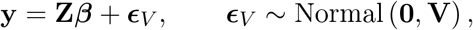

where the design matrix **Z** = (**1, x**_*i*_, …) contains the intercept, genotype and now additional covariates, and ***β*** = (*α, β*_*i*_, …) are their coefficients. The generalized least squares coefficients estimate, used by GCTA and other LMMs, is

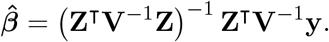

Now suppose **V** corresponds to some kinship matrix **Φ**^*T*^ while **V**′corresponds to **Φ**^*T*^′= *F* ^WG^(**Φ**^*T*^), and **V**′is also invertible. Our strategy involves repeated application of the Sherman-Morrison formula for calculating inverses of matrices after a rank-1 update, which for a symmetric update of a matrix **A** with a vector **z** and a scalar *b* takes the form (Sherman and Morrison, 1950)

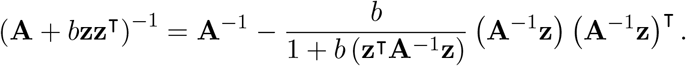

Since *F* ^WG^(**Φ**^*T*^) is a rank-1 update of **Φ**^*T*^ by Eq. (6), then **V**^*t*^ is also a rank-1 update of **V**:

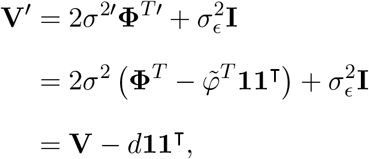

where 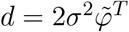 and we used 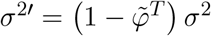. Therefore,

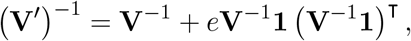

where *e* = *d/ (*1 – *d (***1**^⊤^**V**^*−*1^**1**)). Therefore the following remains a rank-1 update,

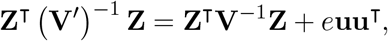

where **u** = **Z**^⊤^**V**^*−*1^**1** is a column vector the length of the number of covariates (including intercept and genotype). Therefore,

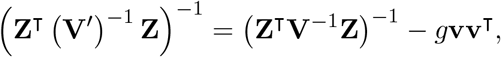

where **v** = (**Z**^⊤^ **V**^*−*1^**Z)**^*−*1^ **u** and *g* = *e/*(1 + *e*(**u**^T^**v**)). Noting that **Z**^T^**V**^*−*1^**1** is the first column of **Z**^⊤^ **V**^*−*1^**Z**, then **v** is the first column of the identity matrix:

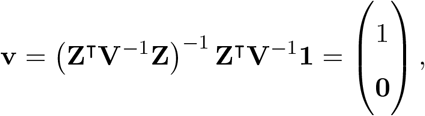

where **0** is a vector the length of the number of covariates minus one (exclude the intercept). As a consequence, **Zv** = **1**, so **u**^⊤^ **v** = **1**^⊤^ **V**^*−*1^**1** and

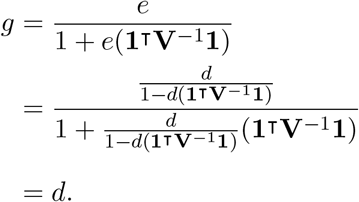

The final step yields the coefficient estimates as a rank-1 update:

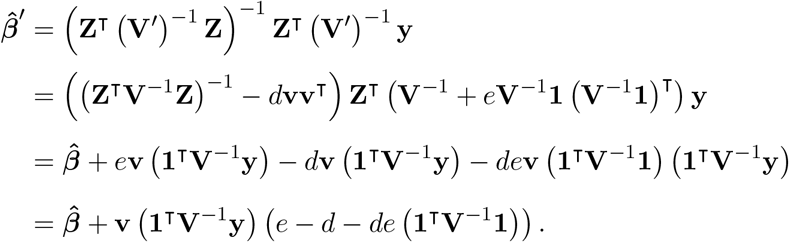

The last factor above vanishes:

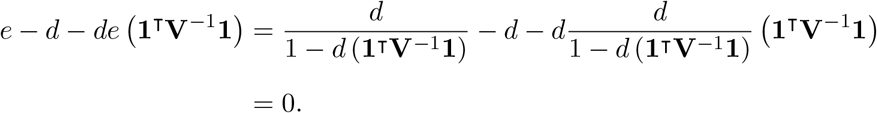

Therefore, 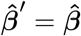, which shows that all fixed effect coefficients, including the intercept, are invariant to using a WG-biased kinship matrix instead of the unbiased one when the coefficients are estimated with generalized least squares.

Furthermore, since the diagonal values of (**Z**^⊤^ (**V**^′^)^*−*1^ **Z)**^*−*1^, which correspond to 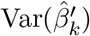 for each *k*, are the same as those of (**Z**^⊤^ **V**^*−*1^**Z)**^*−*1^ except for the first one corresponding to the intercept, then the Wald test statistic of the *k*th covariate coefficients, given by 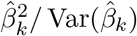, and their p-values, are also the same for *k* ≠ 1 for WG bias as for the unbiased kinship matrix.

## Supplemental figures

**Figure S1:**
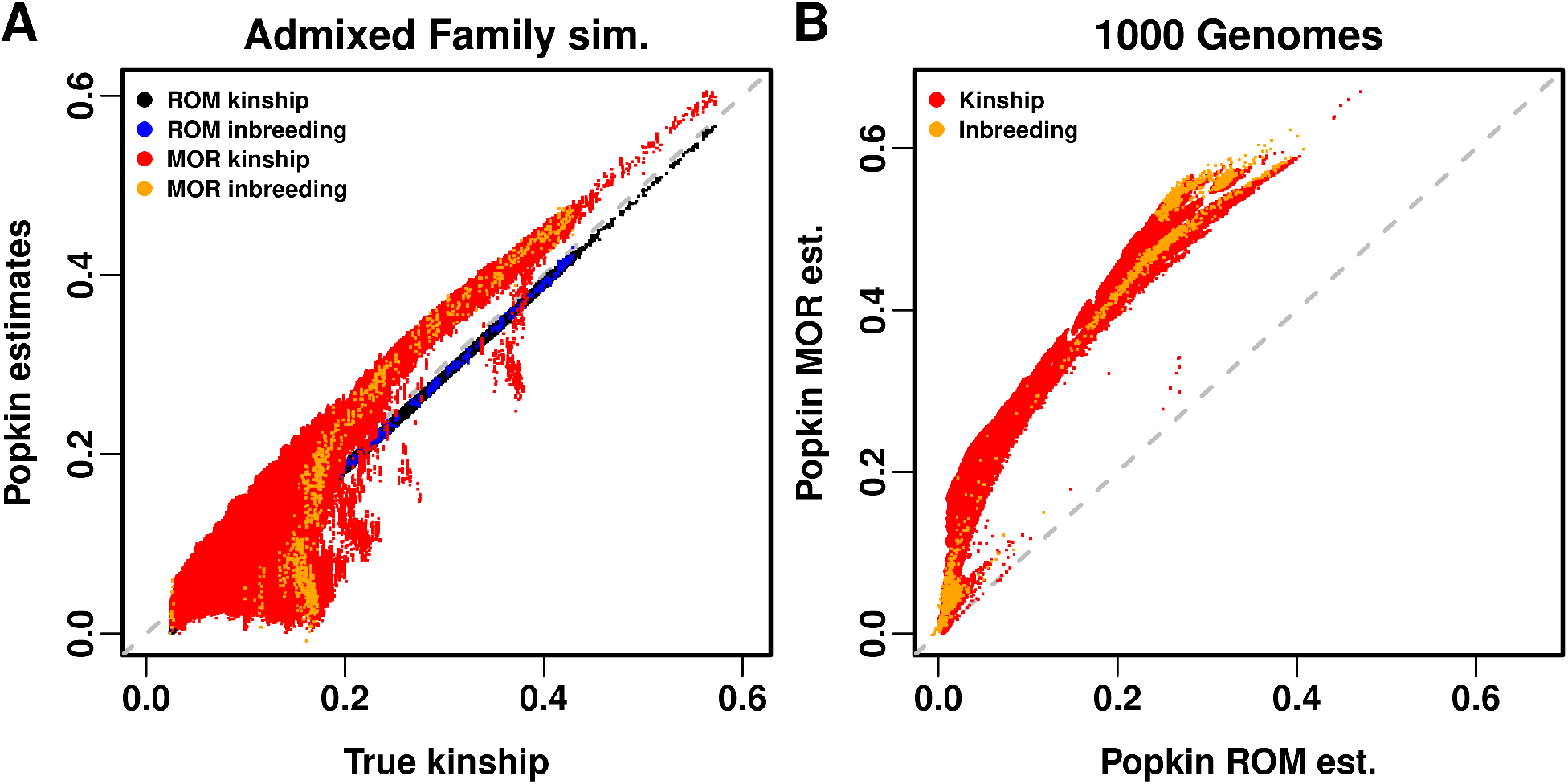
Comparison of popkin ROM and MOR estimates. Kinship (off-diagonal of matrix) and inbreeding (transformed diagonal) are plotted in different colors, which shows that their biases (if any) overlap. **A**. In admixed family simulation, both estimates are compared against true kinship. Popkin ROM has a negligible bias, due to the minimum true kinship of the simulation being slightly larger than zero. Popkin MOR has considerable biases, tending to be upward though not always. **B**. In 1000 Genomes, since true kinship is unknown, popkin ROM takes its place. Popkin MOR biases take on a similar shape as panel A.

**Figure S2:**
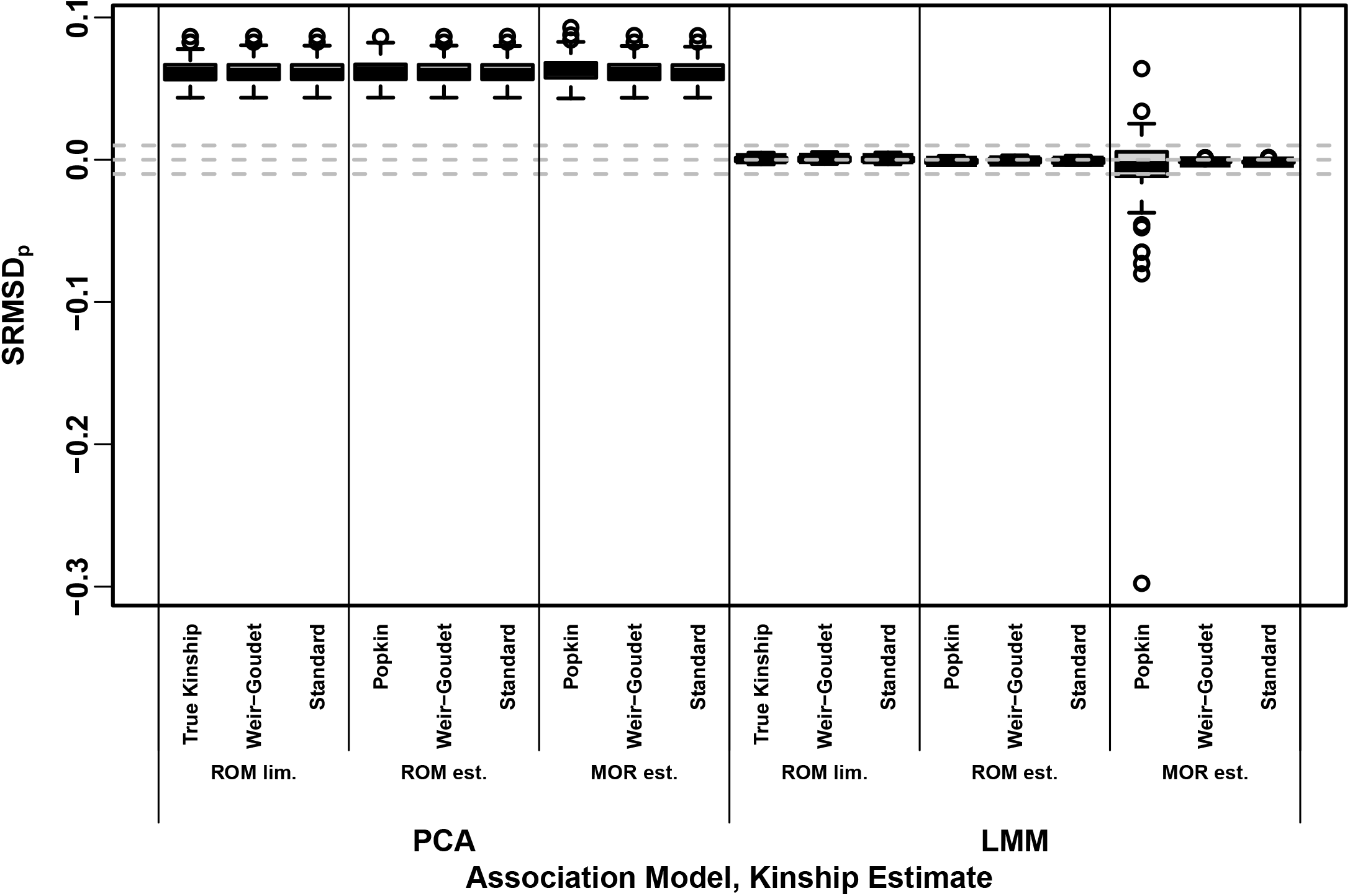
Signed Root Mean Square Deviation of null p-values (SRMSD_*p*_) on the admixed family simulation. Same methods and simulation as Fig. 2, see that for more information. |SRMSD_*p*_| < 0.01 (area between gray dashed lines) is considered calibrated. All PCA runs are miscalibrated by similar amounts, whereas most LMM runs are calibrated with few exceptions.

**Figure S3:**
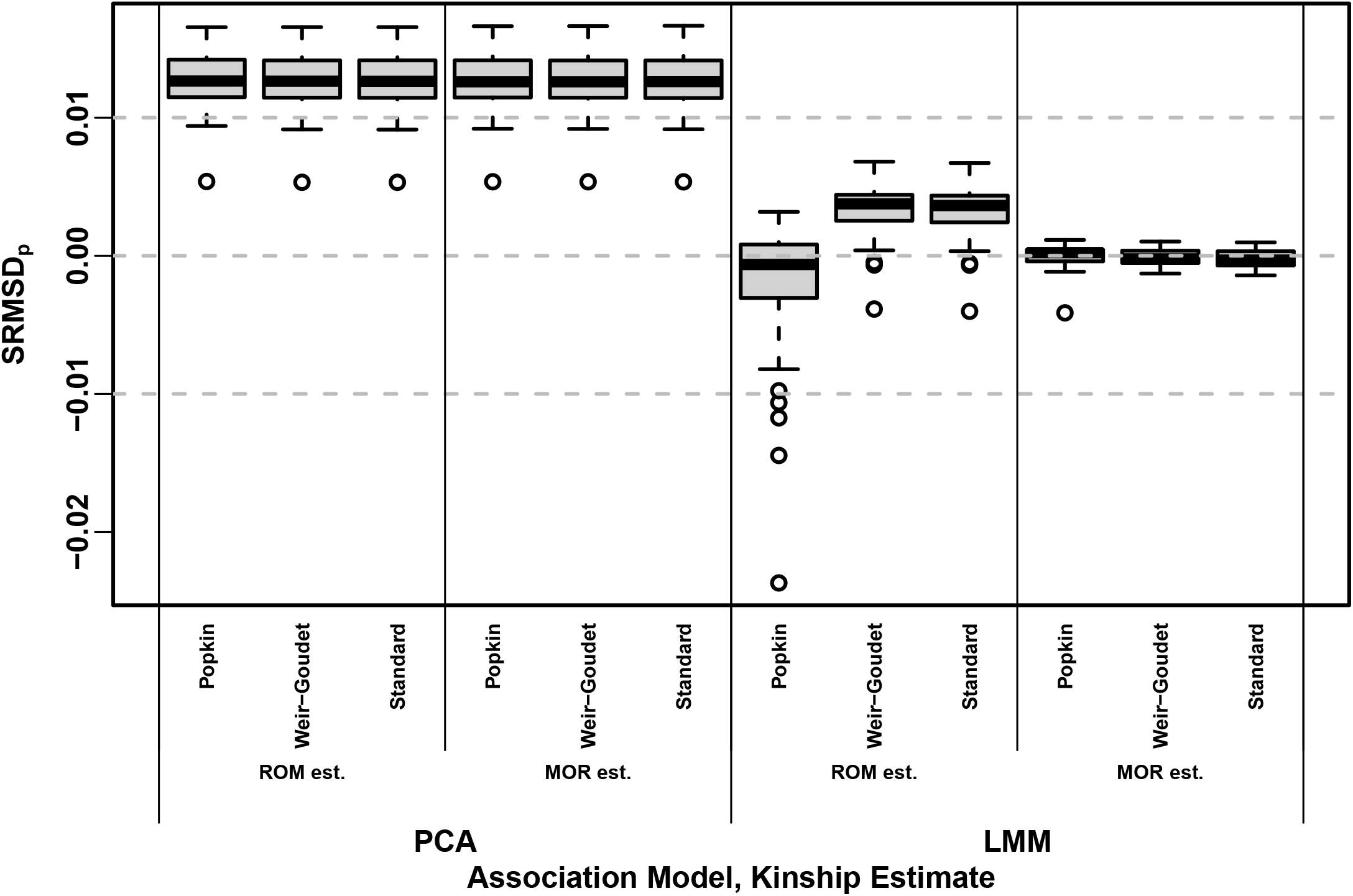
Signed Root Mean Square Deviation of null p-values (SRMSD_*p*_) on 1000 Genomes. Same methods and simulation as Fig. 5, and y-axis statistic and conclusions of Fig. S2, see those for more information.

**Figure S4:**
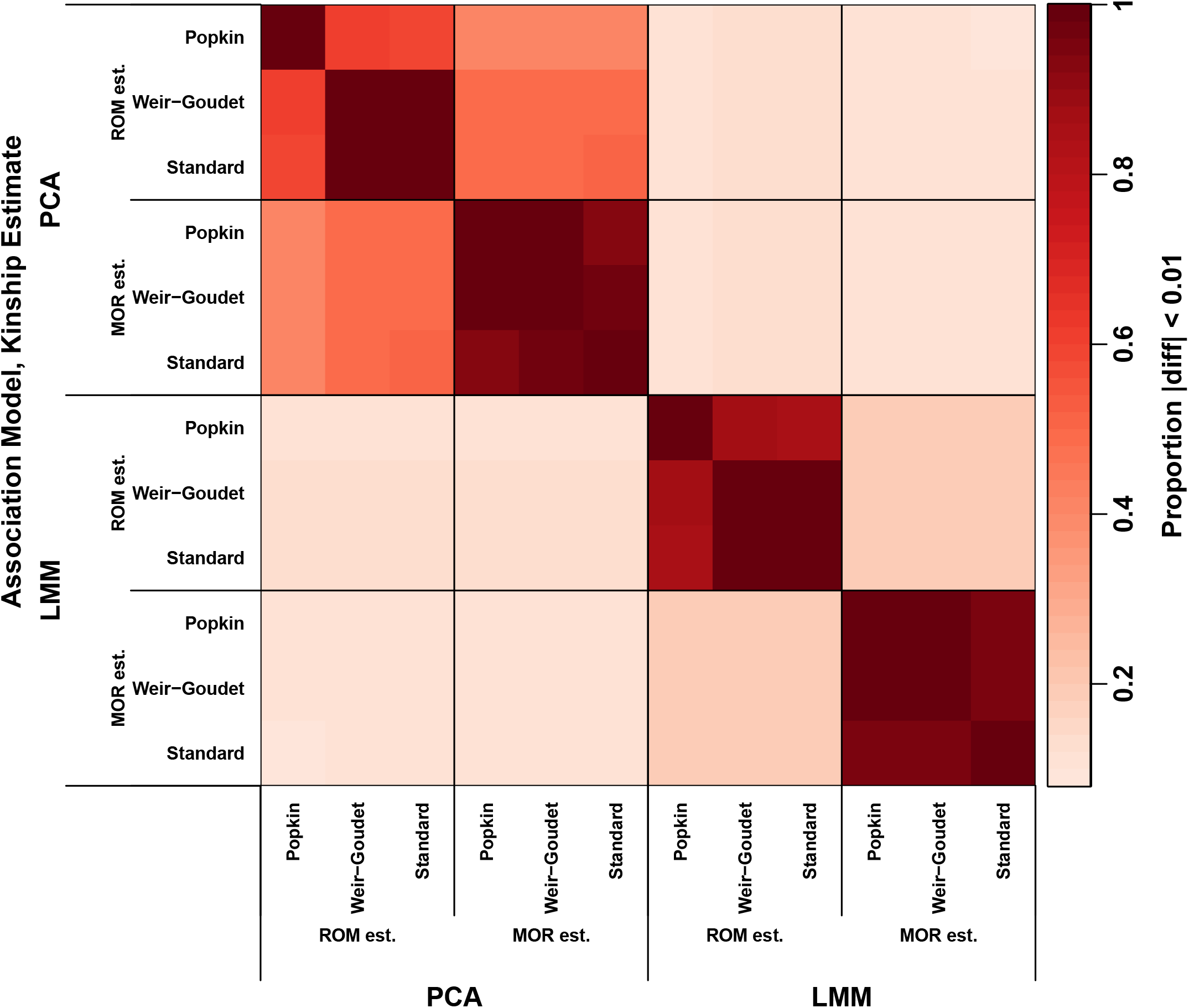
Approximate agreement between p-values on 1000 Genomes. See Fig. 3 for more details.

**Figure S5:**
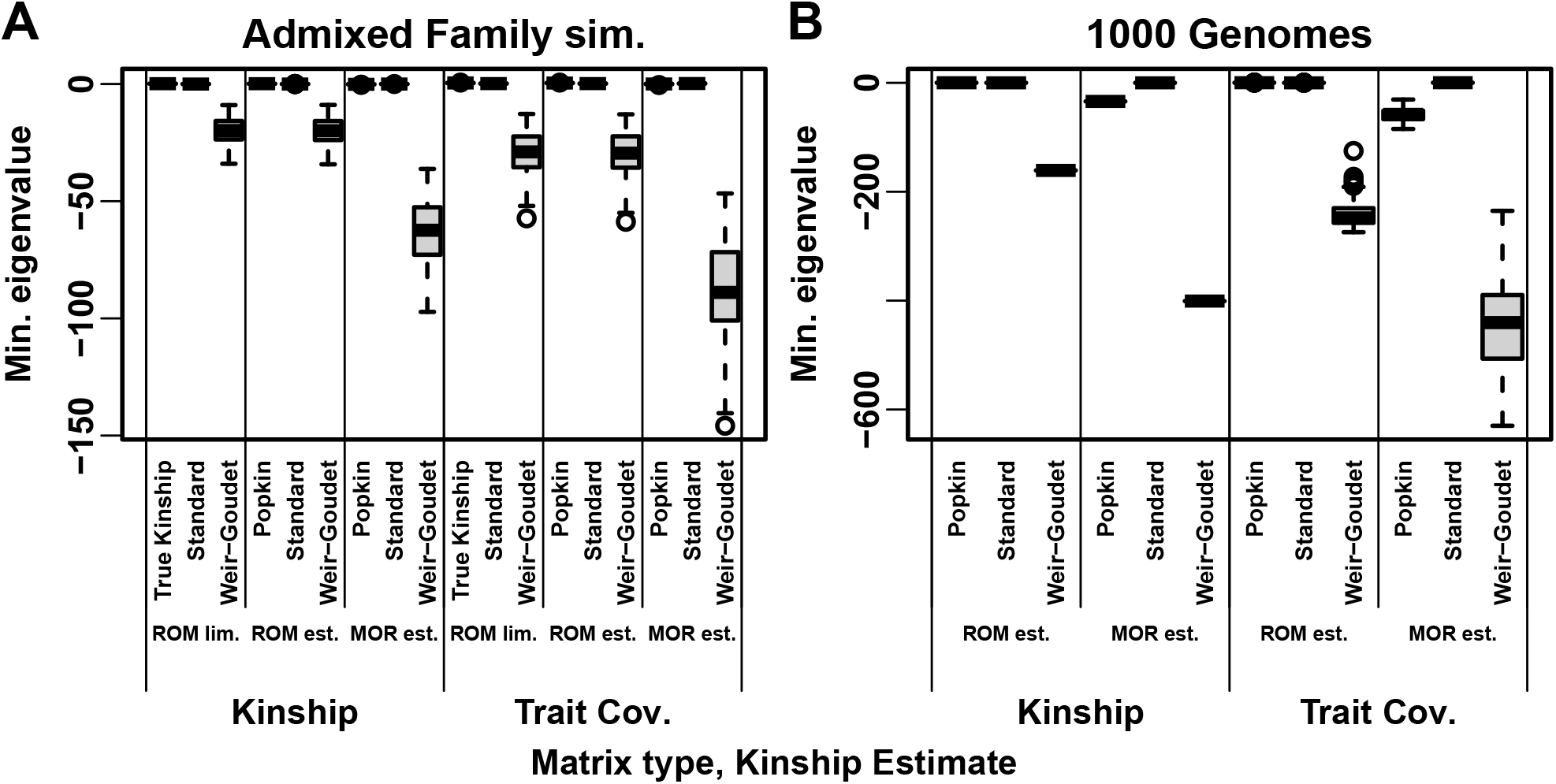
Minimum eigenvalue of kinship and trait covariance (V) matrices. Each distribution is over 100 replicates (1000 Genomes kinship has one value since genotypes are fixed, but **V** varies per replicate). All WG matrices has very large negative eigenvalues, and Popkin MOR has negative eigenvalues as well; in these cases **V** always has negative eigenvalues too.

**Figure S6:**
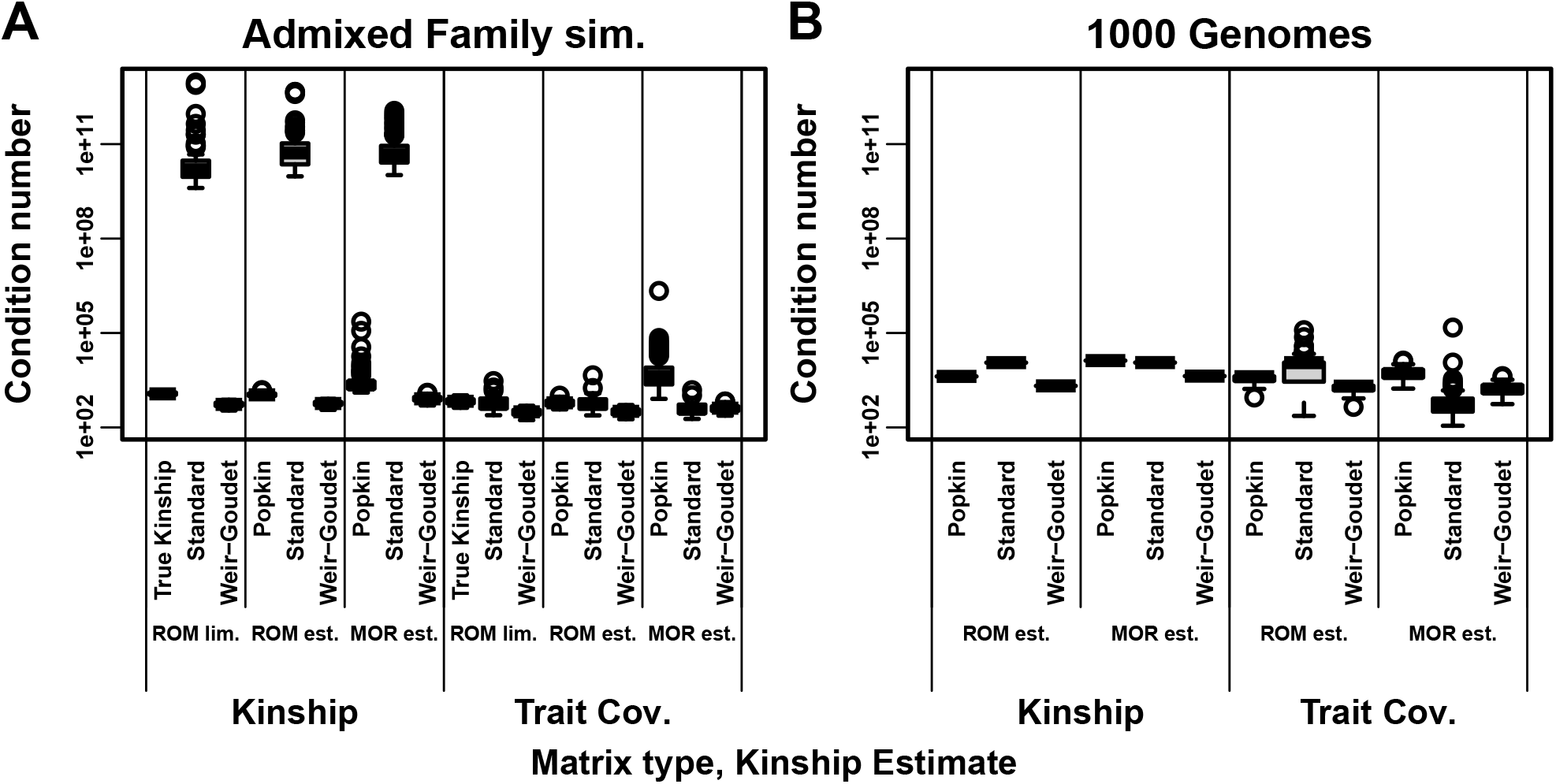
Condition numbers of kinship and trait covariance (V) matrices. Larger condition numbers reflect ill-conditioned problems such as near singularity. Each distribution is over 100 replicates (1000 Genomes kinship has one value since genotypes are fixed, but **V** varies per replicate).

**Figure S7:**
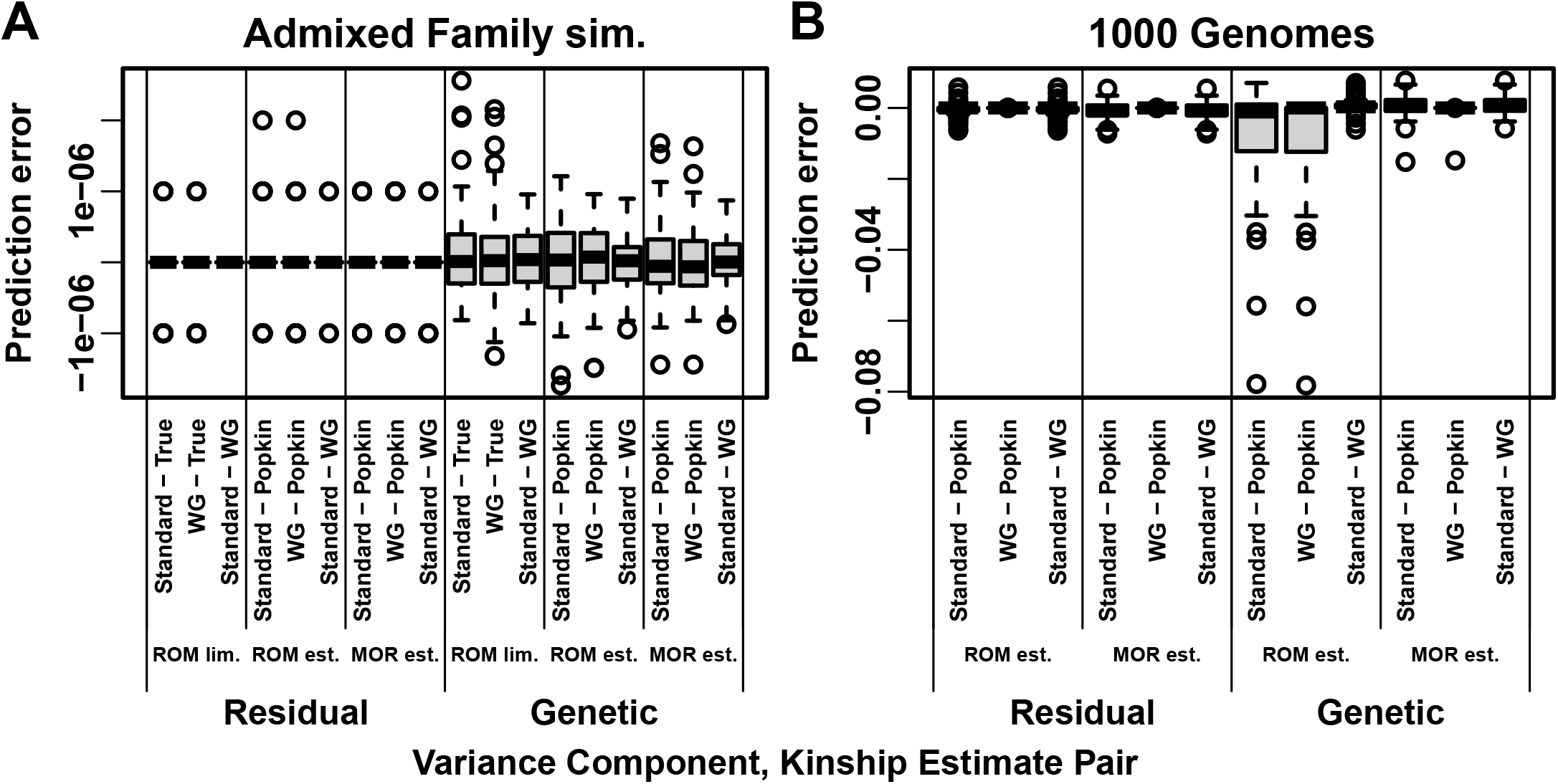
Variance component prediction errors across evaluations. Here we test the predictions that 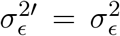 and *σ*^2′^ = *cσ*^2^ in Eq. (16). For Residual, prediction error (y-axis) is 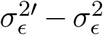 between pairs of estimates as listed. For Genetic, prediction error is *σ*^2′^ − *cσ*^2^: The biased-unbiased pairs use 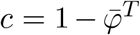 for Standard, 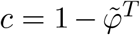 for WG, *σ*^2′^ is their estimate and *σ*^2^ is True or Popkin; The Standard-WG pair uses *σ*^2^ for WG and 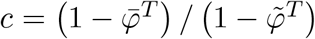. Each distribution is over the 100 replicates of each simulation. **A**. In admixed family simulation, all errors are zero within machine precision. Excess perfect zero residual prediction errors are due to limited precision of GCTA outputs. **B**. In 1000 Genomes, popkin ROM estimates has large errors compared to Standard and WG.

**Figure S8:**
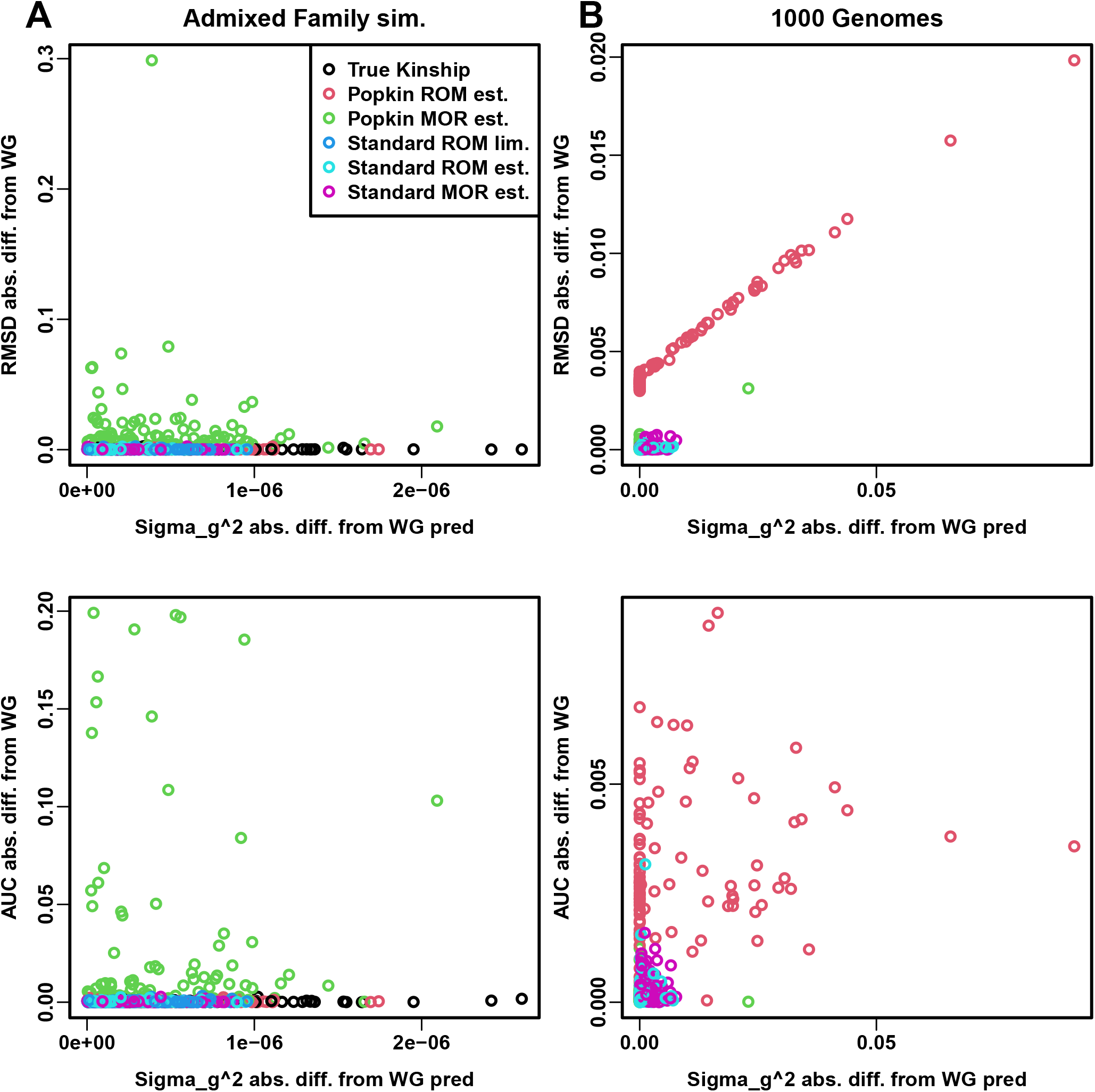
AUC_PR_ and SRMSD_*p*_ prediction errors explained by variance component errors. Genetic variance component (*σ*^2^) absolute error is calculated with the formulas in Fig. S7 using WG as reference since its **V** had the lowest condition numbers (Fig. S6). AUC_PR_ and SRMSD_*p*_ are expected to be the same between WG, Standard, and True or Popkin (within each locus weight type). **A**. Large errors in the admixed family simulation are not explained by high *σ*^2^ error. **B**. Smaller popkin ROM prediction errors in 1000 Genomes are explained by high *σ*^2^ error.

**Figure S9:**
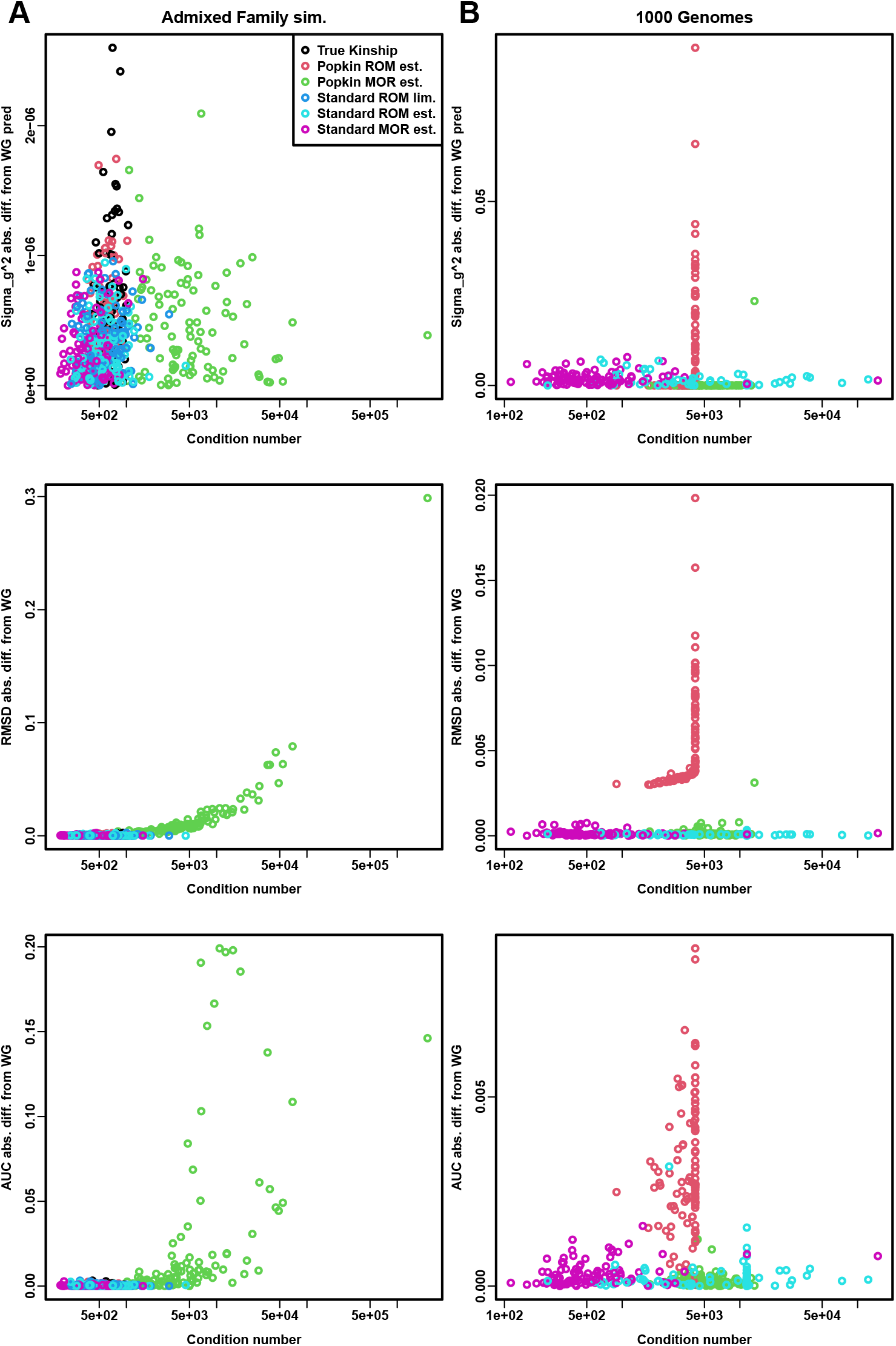
AUC_PR_ and SRMSD_*p*_ prediction errors explained by the condition number of V. AUC_PR_ and SRMSD_*p*_ are expected to be the same between WG, Standard, and True or Popkin (within each locus weight type). WG was used as reference since its **V** had the lowest condition numbers (Fig. S6). **A**. The large popkin MOR prediction errors (AUC_PR_, SRMSD_*p*_, but not *σ*^2^) in the admixed family simulation are explained by the condition number of **V. B**. Smaller errors in 1000 Genomes are not explained by the condition number of **V**.

## Notes

### Competing Interest Statement

The authors have declared no competing interest.

